# Avalanche criticality in individuals, fluid intelligence and working memory

**DOI:** 10.1101/2020.08.24.260588

**Authors:** Longzhou Xu, Lianchun Yu, Jianfeng Feng

## Abstract

The critical brain hypothesis suggests that efficient neural computation can be realized through dynamics of the brain characterized by scale-free avalanche activities. However, the relation between human cognitive performance and the avalanche criticality in large-scale brain networks remains unclear. In this study, we analyzed the mean synchronization and synchronization entropy of blood oxygenation level signals from resting-state fMRI. We found that the scale-free avalanche activity was associated with intermediate synchronization and maximal synchronization entropy. The complexity of functional connectivity, as well as structure-function coupling, is maximized by criticality, which is consistent with theoretical predictions. We observed order-disorder phase transitions in resting-state brain dynamics and found that there were longer times spent in the subcritical regime. These results support the hypothesis that large-scale brain networks lie in the vicinity of a critical point. Finally, we showed evidence that the neural dynamics of human participants with higher fluid intelligence and working memory scores are closer to criticality. We identified brain regions whose critical dynamics showed significant positive correlations with fluid intelligence performance, and found these regions were located in the prefrontal cortex and inferior parietal cortex, which are believed to be important nodes of brain networks underlying human intelligence. Our results reveal the role that avalanche criticality plays in cognitive performance, and provide a simple method to identify the critical point and map cortical states on a spectrum of neural dynamics, ranging from subcriticality to supercriticality.

## Introduction

The critical brain hypothesis states that the brain operates in close vicinity to a critical point that lies between order and disorder. This is characterized by a power-law form of the event size distribution (Cocchi, Gollo, Zalesky, & Breakspear, 2017; Hesse & Gross, 2014). This hypothesis is supported by a set of observations of power-law scaling in many different neural systems using various approaches (J. M. Beggs & Plenz, 2003; Gal & Marom, 2013; Meisel, Olbrich, Shriki, & Achermann, 2013; Plenz, 2012; Shriki et al., 2013; Solovey, Miller, Ojemann, Magnasco, & Cecchi, 2012; Enzo Tagliazucchi, Balenzuela, Fraiman, & Chialvo, 2012). Arguments in favour of this hypothesis have been strengthened by advantages in information transmission, information storage, and dynamic range, in neural systems operating near criticality (Shew, Yang, Petermann, Roy, & Plenz, 2009; Shew, Yang, Yu, Roy, & Plenz, 2011; Yang, Shew, Roy, & Plenz, 2012), with evidence arising in both theoretical and experimental work (Shew & Plenz, 2012). Meanwhile, this hypothesis still faces challenges from several perspectives (J. Beggs & Timme, 2012). For example, computational studies suggested that power laws may emerge from simple stochastic processes or noncritical neuronal systems (Touboul & Destexhe, 2010), so power laws alone are prerequisite but not sufficient evidence for criticality. Meanwhile, it has been asked: “If the brain is critical, what is the phase transition (Fontenele et al., 2019)?” Indeed, the observation of power law avalanche activity along with a phase transition between order and disorder would be more persuasive for criticality. Furthermore, though previous studies have associated supercriticality with reduced consciousness (Meisel et al., 2013; Scott et al., 2014), near-critical dynamics with rest (Priesemann et al., 2014), and subcriticality with focused cognitive states (Fagerholm et al., 2015), there remains a gap between the specific brain state and efficient information processing endowed by criticality as predicted by theory (He, 2011). To fully understand the functional roles of critical and non-critical dynamics, more research is required to relate brain states and cognitive performance to neural dynamics that lie on a spectrum, ranging from subcriticality to supercriticality. To obtain a deeper understanding of this phenomenon, it is necessary to develop data analysis methods to represent this phase spectrum with high resolution and characterize the subsequent reorganization of brains with the transition in this spectrum (Fontenele et al., 2019).

With advances in brain imaging techniques such as functional magnetic resonance imaging (fMRI), the critical brain hypothesis has found roles in interpreting fundamental properties of large-scale brain networks in the context of structure-dynamics-function relationships (Karahanoğlu & Van De Ville, 2017; Lee et al., 2019). For example, it has been shown that structural connections of brains are mostly reflected in functional networks, and this structure-function coupling is disrupted when brains move away from criticality during anesthesia (Enzo Tagliazucchi et al., 2016). Another application of criticality is to explain the dynamic basis of brain complexity (Popiel, 2020; E. Tagliazucchi & Chialvo, 2013; Timme et al., 2016). In particular, functional connectivity (FC) complexity, which is an umbrella term describing the variability, diversity, or flexibility of functional connections in brain networks, has been associated with cognitive performance from many perspectives, such as high-order cognition, aging, and cognitive impairment in brain disorders (Ahmadlou, Adeli, Bajo, & Adeli, 2014; Anokhin, Birbaumer, Lutzenberger, Nikolaev, & Vogel, 1996; Omidvarnia, Zalesky, Van De Ville, Jackson, & Pedersen, 2019; Smyser et al., 2016; B. Wang et al., 2017). Studies have suggested that the FC complexity may possibly be at its maximum at the critical point, while the FC capacity arises from special topological properties of the structural network, such as hierarchical modular organization (Song et al., 2019; R. Wang et al., 2019).

To validate these applications, both computer modeling methods and experimental data analysis methods were used. Computer modeling utilizes structural imaging data to model large-scale brain dynamics and functional networks (Deco, Jirsa, & McIntosh, 2011; Nakagawa, Jirsa, Spiegler, McIntosh, & Deco, 2013). However, there is still disagreement on which type of phase transition should be adopted for large-scale brain networks, e.g., first-order discontinuous vs. second-order continuous phase transitions and edge of chaos criticality vs. avalanche criticality (Kanders, Lorimer, & Stoop, 2017; Scarpetta, Apicella, Minati, & de Candia, 2018). Experimental studies usually take advantage of dynamic changes caused by interventions, such as deprived sleep, anesthesia, or brain diseases, to show deviations from criticality and subsequent reorganization of FC networks (Hobbs, Smith, & Beggs, 2010; Meisel et al., 2013; Meisel, Storch, Hallmeyer-Elgner, Bullmore, & Gross, 2012; Enzo Tagliazucchi et al., 2016). However, deviations caused by these interventions are usually unidirectional, either in the sub- or supercritical directions. Furthermore, it is not clear whether deviations from criticality caused by different intervention methods follow an identical phase transition trajectory. Recent studies have proposed the concept of a “critical line” instead of a “critical point” and suggested that multiple phase transition trajectories may exist (Kanders, Lee, Hong, Nam, & Stoop, 2020). Therefore, the successful retrieval of the phase transition trajectory from the large-scale brain networks will not only help to answer key questions regarding what the phase transition is if the brain is critical but also have important implications in brain functional imaging and large-scale brain modeling.

In this study we applied a large number of criticality-related metrics to investigate the spatio-temporal dynamics of large scale brain networks as well as their variation in individuals, and explored associations with three different cognitive abilities in a sample of 295 healthy young adults from the Human Connectome Project (HCP) 1200-subject release (Van Essen et al., 2013). We retrieved an inverted-U curve by mapping individuals’ brains onto the phase plane between the mean synchronization (MS) and synchronization entropy (SE) of blood oxygenation level-dependent (BOLD) signals. We then performed the classic avalanche criticality analysis method on these data (J. M. Beggs & Plenz, 2003; Enzo Tagliazucchi et al., 2012), and found that subjects who were closest to the critical point exhibited moderate mean and maximal variability in synchrony of BOLD signals (i.e., located around the tipping point of the inverted-U curve). This is consistent with previous findings that the neural systems operate neither at the synchronous nor the asynchronous ends of the spectrum, but rather near the critical point between them (Fontenele et al., 2019). We also showed that the large individual variation around the critical point could be the cause of the mismatch between group-level data analysis and theoretical prediction. However, this individual variation also gave us a chance to examine previous conjectures on criticality in large-scale brain networks. And we indeed found that both FC complexity and structure-function coupling were maximized around the criticality. We also utilized a sliding window approach to observe “instantaneous” phase transition occurring in individual brains. We found that brains persisting in the subcritical regime exhibited longer dwell times than those in other regimes. Finally, we found the critical dynamics were associated with high scores in fluid intelligence and working memory tests, but not with crystallized intelligence scores. Additionally, the critical dynamics in the frontal cortex, superior parietal lobule, angular gyrus and supramarginal gyrus et al, which were believed as vital regions in the networks of Parieto-Frontal Integration Theory (P-FIT) for intelligence (Jung & Haier, 2007), exhibited significant correlations with fluid intelligence performance.

## Material and methods

### Data acquisition and preprocessing

#### fMRI data acquisition and preprocessing

We used rfMRI data from the Human Connectome Project (HCP) 1200-subject release (Van Essen et al., 2013). Each subject underwent two sessions of rfMRI on separate days, each session with two separate 14 min 24s acquisitions generating 1200 volumes on a customized Siemens 3T Skyra scanner (TR = 720 ms, TE = 33 ms, flip angle = 52 °, voxel size = 2 mm isotropic, 72 slices, FOV = 208 × 180mm, matrix = 104 × 90 mm, multiband accelaration factor = 8, echo spacing = 0.58 ms). The rfMRI data used for our analysis were processed according to the HCP minimal preprocessing pipeline (Glasser et al., 2016; Glasser et al., 2013) and denoising procedures. The denoising procedure pairs the independent component analysis with the FSL tool FIX to remove non-neural spatiotemporal components (Smith et al., 2015). And as a part of cleanup, HCP used 24 confound time series derived from the motion estimation (the 6 rigid-body parameter time series, their backwards-look temporal derivatives, plus all 12 resulting regressors squared). Note that the global component of the fMRI fluctuations measured during the resting state is tightly coupled with the underlying neural activity, and the use of global signal regression as a pre-processing step in resting-state fMRI analyses remains controversial and is not universally recommended (Liu, Nalci, & Falahpour, 2017). Therefore, the global whole-brain signal was not removed in this work. We used the left-to-right acquisitions from the first resting-state dataset (i.e., resting state fMRI 1 FIX-denoised package).

The first 324 subjects in the dataset entered into our study, and we excluded 29 subjects for missing data. This left us with 295 subjects for further analysis, and 162 of them were females. All the participants were between the ages of 22-36, 58 were between the ages of 22-25, 130 were between the ages of 26-30, 104 were between the ages of 31-35, and 3 were 36 years old.

For further analysis, the whole cortex was parcellated into 96 regions using the Harvard-Oxford atlas (Makris et al., 2006), and the details are provided at https://identifiers.org/neurovault.image:1699, and from each region the time series averaged across the voxels were extracted and Z-normalized to construct region of interest (ROI) signals (atlas_96_ signals). To test the results for different parcellation, we also used the Human Brainnetome atlas that comprised 246 regions (Fan et al., 2016), and Zalesky atlas that comprised 1024 regions (Zalesky et al., 2010), the resulting signals were termed as atlas_246_ and atlas_1024_ signals, respectively.

#### Diffusion tensor imaging (DTI) data acquisition and preprocessing

The diffusion MRI images used in this study were also from the HCP 1200-subject release (Sotiropoulos et al., 2013). Briefly, the diffusion data were collected using a single-shot, single refocusing spin-echo, echo-planar imaging sequence (TR = 5520 ms, TE = 89.5 ms, flip angle = 78 °, FOV = 210 × 180 mm, matrix = 168 × 144 mm, voxel size = 1.25 mm istropic, slices = 111, multiband acceleration factor = 3, echo spacing = 0.78 ms). Three gradient tables of 90 diffusion-weighted directions and six b=0 images, each were collected with right-to-left and left-to-right phase encoding polarities for each of the three diffusion weightings (b=1000, 2000, and 3000 s/mm^2^). All diffusion data were preprocessed with the HCP diffusion pipeline updated with EDDY 5.0.10 (Sotiropoulos et al., 2013), and the details are provided at https://www.humanconnectome.org. In this study, from the 295 selected subjects, only 284 subjects were entered into our DTI data analysis because 11 of them were missing the corresponding DTI data.

#### Cognition measures

We examined associations between cognitive ability and critical dynamics conducted in our rfMRI analysis. Three relevant behavioral tasks were used as a measure of cognitive ability, including fluid intelligence, working memory, and crystallized intelligence. The same 295 subjects were also entered into our cognitive ability analysis, but for the fluid intelligence analysis, 5 subjects were excluded due to missing their intelligence scores, or information about age and education.

The fluid intelligence scores in the HCP data release were measured using the number of correct responses on form A of the Penn Matrix Reasoning Test (PMAT, mean = 17.0034, standard deviation (SD) = 4.9106, range = 4 − 24), which had 24 items and 3 bonus items, using nonverbal visual geometric designs with pieces to assess reasoning abilities that can be administered in under 10 min (Barch et al., 2013; Hearne, Mattingley, & Cocchi, 2016). The PMAT (Bilker et al., 2012) is an abbreviated version of Raven’s Standard Progressive Matrices test (Wendelken, Nakhabenko, Donohue, Carter, & Bunge, 2007), which comprises 60 items.

Crystallized intelligence was measured using the picture vocabulary test (Picture vocabulary, mean=116.8205, SD=10.1977, range=92.3914-153.0889) from the National Institutes of Health (NIH) toolbox (Barch et al., 2013; Hearne et al., 2016). This measure of receptive vocabulary was administered in a computer-adaptive testing (CAT) format. The participant was presented with four pictures and heard an audio recording saying a word, and was instructed to select the picture that most closely showed the meaning of the word. Because the test used a variable length CAT with a maximum of twenty-five items, some participants had fewer items, and the presented words depended on the participant’s performance.

Working memory was assessed using the List Sorting Working Memory test (List sorting, mean=111.2075, SD=12.0946, range=84.63-144.50) from the NIH Toolbox (Barch et al., 2013), in which the participants were required to sequence sets of visually and a small number of orally presented stimuli in size order from smallest to biggest. Pictures of different foods and animals were displayed with both a sound clip and a written test that names them, and involved two different conditions. In the 1-list condition, participants ordered a series of objects, either food or animals, but in the 2-list condition, participants were presented with both animal and food lists and asked to order each list by increasing size. The number of list items increased in subsequent trials, and the task was discontinued after 2 consecutive incorrect trials.

### Data analysis methods

#### Synchrony and variability in synchrony

We measured the mean and variability in synchronization with a previously described approach (Meisel et al., 2013; Yang et al., 2012). First, we obtained the phase trace *θ*(*t*) from the signal *F_j_*(*t*) using its Hilbert transform *H*[*F_j_*(*t*)]:

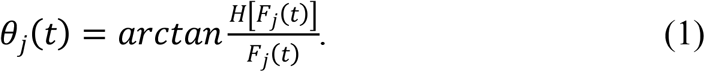

Next, we calculated the Kuramoto order parameter as follows:

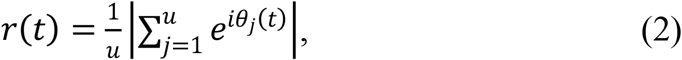

in which u is the number of ROIs in global network analysis, or the number of voxels in a particular region in regional analysis. The Kuramoto order parameter *r*(*t*) was used as a time-dependent measure of phase synchrony of a system. The MS of a time period was calculated as

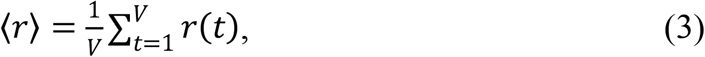

where *V* is the length of the time period. In this study, we calculated static MS of the entire scan period with *V* = 1200 time points. We derived the entropy of *r*(*t*) as the measure of variability in synchronization (synchronization entropy, SE):

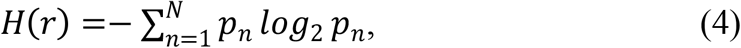

where *p_n_* is the probability that *r*(*t*) falls into a bin between *min* (*r*(*t*)) ≤ *b_n_* < *r*(*t*) < *b*_*n*+1_ ≤ *max* (*r*(*t*)). In this study, we chose the number of bins *N* = 30, and the robustness of our results was also tested within an interval between 5 and 100.

#### Avalanche analysis

In our avalanche analysis, the ROI signals were reduced to a spatiotemporal point process by detecting the suprathreshold peak positions intermediate between two above-threshold time points, as shown in the example in Fig. 1A. By binning the binary sequences with appropriate time resolution (time bin), we obtained a spatial pattern of active ROIs within consecutive time bins. An avalanche was defined as a series of consecutively active bins, which were led and followed by blank bins without activation. The size S and duration T of the avalanches were then defined as the total number of activations and total number of time bins during this avalanche, respectively (J. M. Beggs & Plenz, 2003; Enzo Tagliazucchi et al., 2012).

**Figure 1.**
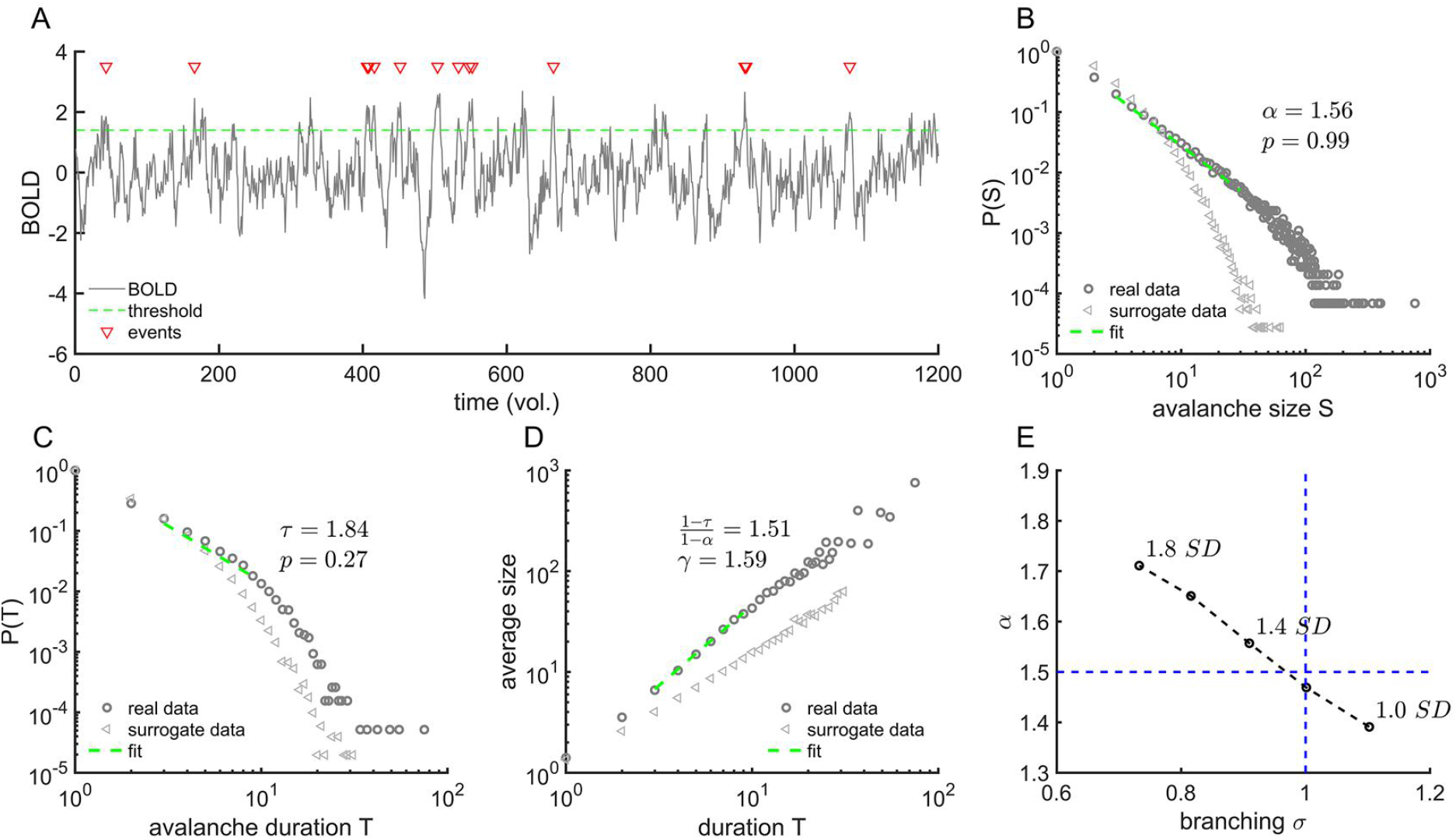
Avalanche statistics obtained from group-level analysis. **A.** Example of a point process (red triangles) extracted from one normalized ROI BOLD signal. **B.** The probability distributions of group-aggregated avalanche sizes for the threshold 1.4 SD and the time bin width of 1 volume (vol.) in the fMRI data. The distributions are well approximated by power law with an exponent of *α* = 1.56 with Clauset’s test *p* = 0.99, corresponding to *κ* = 0.9912 (histogram bins=40) and branching parameter *σ* = 0.9097. **C.** The distribution of avalanche durations can be fitted well by a power law with an exponent of *τ* = 1.84 under the condition described in **B**, with Clauset’s test *p* = 0.3. **D.** There is a relation between the sizes and duration of the avalanches with a positive index *γ* = 1.59, which is close to 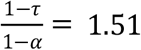. In **B-D**, the grey open triangles were calculated from the surrogate data. **E.** The branching ratio and power-law scaling exponents *α* of avalanche sizes for different thresholds used to define the point process.

If a system operates near a critical point, the size distribution (*P*(*S*)), duration distribution (*P*(*T*)), and average size for a given duration (〈*S*〉(*T*)) should be fitted into power laws:

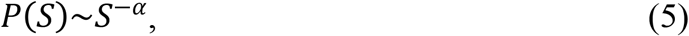

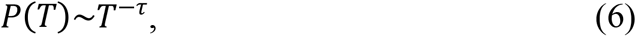

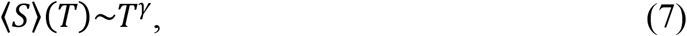

where *α, τ*, and *γ* are critical exponents of the system (Friedman et al., 2012; Sethna, Dahmen, & Myers, 2001). Furthermore, the following scaling relation was proposed as an important evaluation of the criticality (Fontenele et al., 2019; Friedman et al., 2012), namely,

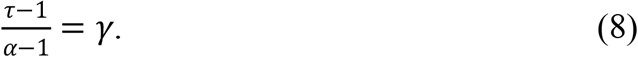

In this study, we defined

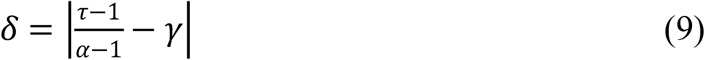

to measure the distances of the systems from the critical point, so the smaller *δ* is, the closer the systems are to the critical point.

The scaling exponents governing the power-law distribution were estimated using the maximum likelihood estimator (MLE) (Clauset, Shalizi, & Newman, 2009; Marshall et al., 2016). Briefly, the MLE procedure supposes that the empirical data sample is from a power-law function in the range (*x_min_, x_max_*), with probability density 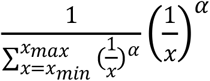 (Fontenele et al., 2019; Marshall et al., 2016). We estimated critical exponents *α* and *τ* by maximizing the likelihood function and via a lattice search algorithm (Marshall et al., 2016). We then used Clauset’s goodness-of-fit test to quantify the plausibility of fits (Clauset et al., 2009; Deluca & Corral, 2013; Marshall et al., 2016). We used a power-law model to produce data sets over the fit range and compared the Kolmogorov–Smirnov (KS) statistics between (1) the real data and the fit against (2) the model data and the fit. If the real data produced a KS-statistic that was less than the KS-statistic found for at least 10% of the power-law models (i.e., p ≥ 0.1), we accepted the data as being fit by the truncated power law because the fluctuations of the real data from the power law were similar in the KS sense to random fluctuations in a perfect power-law model.

#### Surrogate data

To assess the statistical significance of the avalanche analysis results and MS-SE relationship, we generated comparable surrogate data and applied the analyses above to these data. Phase-shuffling is often used in hypothesis testing for avalanche size distribution (Gireesh & Plenz, 2008; Shriki et al., 2013). Phase-shuffling disrupts temporal as well as spatial correlations in multichannel time series.

Herein phase shuffling was done on the atlas_96_ signals. The phase randomization procedures were as follows (Prichard & Theiler, 1994): (1) the discrete Fourier transformation was taken to of each subject; (2) rotating the phase at each frequency by an independent random variable that was uniformly chosen in the range [0,2*π*]. Crucially, the different time series were rotated by the different phases to randomize the phase information; (3) the inverse discrete Fourier transformation was applied to these time series to yield surrogate data.

#### Branching parameter

The branching parameter *σ*, which is defined as the average number of subsequent events that a single preceding event in an avalanche triggers, is a convenient measure to identify criticality (J. M. Beggs & Plenz, 2003). In theory, the system is critical for σ = 1 and sub-(super) critical for σ < 1 (σ > 1). In this study, σ was calculated as

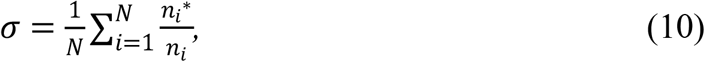

where *n_i_* is the number of ancestors, *n_i_\** is the number of descendants in the next time bin, and *N* is the total number of time bins with activations.

#### Definition of kappa

A nonparametric measure, *κ*, for neuronal avalanches was introduced by Shew and his colleagues (Shew et al., 2009). It quantifies the difference between an experimental cumulative density function (CDF) of the avalanche size, *F*(*β_k_*), and the theoretical reference CDF, *F^NA^*(*β_k_*), which is a power-law function with theoretical expected exponent *α* = 1.5:

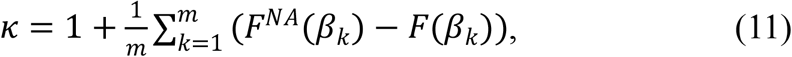

where *β_k_* are avalanche sizes logarithmically spaced between the minimum and maximum observed avalanche sizes, and *m* is the number of histogram bins. The unit value of *κ* is characteristic of the system in a critical state, whereas values below and above 1 suggest sub- and supercritical states, respectively.

#### Functional and structure connectivity matrix

We constructed an FC matrix from atlas_96_ signals by computing the Pearson correlation *C_ij_* between ROI *i* and ROI *j*, and the mean FC strength 〈*FC*〉 was obtained by

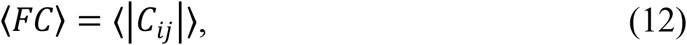

where |·| means the absolute value.

The structure connectivity (SC) matrix was constructed using DSI Studio (http://dsi-studio.labsolver.org) from DTI data. The DTI data were reconstructed in the Montreal Neurological Institute (MNI) space using q-space diffeomorphic reconstruction (F.-C. Yeh & Tseng, 2011) to obtain the spin distribution function (F. Yeh, Wedeen, & Tseng, 2010). A diffusion sampling length ratio of 1.25 was used. The restricted diffusion was quantified using restricted diffusion imaging (F.-C. Yeh, Liu, Hitchens, & Wu, 2017), and a deterministic fiber tracking algorithm (F.-C. Yeh, Verstynen, Wang, Fernández-Miranda, & Tseng, 2013) was used to obtain one million fibers with whole-brain seeding. The angular threshold was randomly selected from 15 degrees to 90 degrees. The step size was randomly selected from 0.1 voxels to 3 voxels. The anisotropy threshold was automatically determined by DSI Studio. The fiber trajectories were smoothed by averaging the propagation direction with a percentage of the previous direction. The percentage was randomly selected from 0% to 95%. Tracks with a length shorter than 5 or longer than 300 mm were discarded. The SC matrix was calculated by using the count of the connecting tracks using 96-region Harvard-Oxford atlas.

#### Functional connectivity entropy

The FC entropy *H*(*FC*) is calculated by

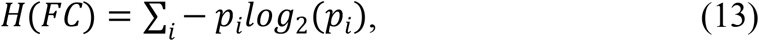

where *p_i_* is the probability distribution of |*C_ij_*|, i.e., ∑_*i*_*p_i_* = 1 (Yao et al., 2013). In the calculation, the probability distribution was obtained by discretizing the interval (0, 1) into 30 bins.

#### Functional connectivity diversity

The functional diversity (*D*(*FC*)) of the FC matrix is measured by the similarity of the distribution to the uniform distribution (R. Wang et al., 2019):

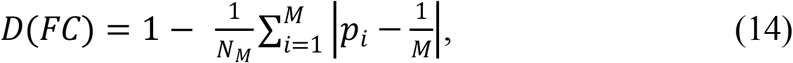

where 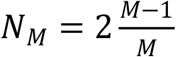 is a normalization factor, *D*(*FC*) is in the range [0, 1], and *P_i_* is the probability distribution of |*C_ij_*|, which was obtained by discretizing the interval (0, 1) into M bins (*M* = 30 in this work). For completely asynchronous or synchronized states, the correlation values fall into one bin at 0 or 1, where *D*(*FC*) = 0 reflects the simple dynamic interaction pattern. In an extreme case where all types of FC equivalently exist, *p_i_* would ideally follow a uniform distribution (i.e., probability in each bin = 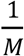) and *D*(*FC*) = 1.

#### Functional connectivity flexibility

To obtain the flexibility of FC in the whole brain, we utilized the sliding window method to calculate connectivity number entropy (*CNE*) for each region (Lei et al., 2020; Song et al., 2019). A non-overlapping sliding window method was applied to the atlas_96_ signals. The choice of window size must be sufficient to yield a stable Pearson’s correlation coefficient within each window, yet small enough to reveal the temporal-dependent variation in FC (Lei et al., 2020; Sakoğlu et al., 2010). We chose a window size in the range of 20-30, corresponding to the number of windows (*n_win_*) in the range of 40-60.

Within each time window, we first acquired the FC matrix via their time series in this window. Then, the binary network matrix was obtained by binarizing the FC matrix with a threshold *THR_FC_*. Subsequently, we calculated the number of regions connected to a particular region *k* (*k* = 1, 2,… 96) in each time window. Therefore, we could obtain *p_i_*, the probability for a particular connection number occurring, where *i* indicated the *i-th* connection number among all possible connection numbers. Then, for the region *k*, *CNE_k_* is a complexity measure (i.e., Shannon entropy) for the disorder in the connection numbers over time:

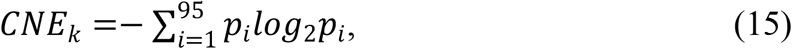

where the summation index runs from 1 to the number of all possible connection numbers.

For each subject, the *CNE* at the whole-brain level was obtained by simply averaging the regional *CNE_k_* values over 96 regions:

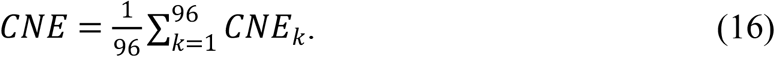

#### Similarity between functional and structural networks

To measure the similarity between functional and structural connection networks, the FC matrices *C_ij_* were thresholded by *THR_FC_* to yield binary adjacency matrices *A_ij_* such that *A_ij_* = 1 if *C_ij_* ≥ *THR_FC_*, and *A_ij_* = 0 otherwise. The parameter *THR_FC_* was chosen to fix link density *ρ_FC_*, which was defined as the ratio of the connections in the network (∑_*i*>*j*_ *A_ij_*) to the total possible number of connections. It is important to fix the link density when comparing networks, as otherwise, differences could arise because the average of the respective *C_ij_* are different (and therefore the number of nonzero entries in *A_ij_*) but not because connections are topologically reorganized across conditions (Enzo Tagliazucchi et al., 2016).

The binary FC networks for each subject were compared with the group-aggregated binary SC network but not with the individual’s SC network to avoid fluctuations in individual SC networks. First, the binary adjacency matrices *B_ij_* of SC matrices were obtained for each subject such that *B_ij_* = 1 if there were tracked fiber links; otherwise, *B_ij_* = 0. Then, the binary adjacency structural connection matrices were summed up and again thresholded by a thresholding value *THR_SC_* to yield a group-aggregated binary SC network. In this way, high *THR_SC_* values would exclude connections that were shared by fewer subjects but preserve connections that were common in most subjects.

To estimate the similarity between the binary FC network of each subject and the group-aggregated binary SC network, we computed the Pearson correlation *R*(*FC* − *SC*) and Hamming distance *HD*(*FC* − *SC*) between these two networks (Enzo Tagliazucchi et al., 2016). Specifically, the Hamming distance is defined as the number of symbol substitutions (in this case 0 or 1) needed to transform one sequence into another and vice versa, and in this case, it is equal to the number of connections that must be rewired to turn the functional network connection into the structural network connection.

#### Dynamic analysis of phase transitions

We used the sliding window approach to capture the time-dependent changes in measures used in this study. In the calculation, for the atlas_96_ signals, the length of the sliding window was set to *V* = 200 (volumes), and the sliding step was set to Δ*n* = 10 (volumes). In each window, we calculated the corresponding dynamic measures, including dynamic MS 〈*r*〉_*n*_, dynamic SE *H*(*r*)_*n*_, and dynamic FC matrix (*C_ij_*)_*n*_. From the dynamic FC matrix (*C_ij_*)_*n*_, we further obtained dynamic FC entropy *H*(*FC*)_*n*_, FC diversity *D*(*FC*)_*n*_, Pearson correlation between FC and SC R(FC − SC)_n_, and Hamming distance *HD*(*FC* − *SC*)_*n*_.

#### Data and code availability

The rfMRI data, DTI data and cognitive data are available from the Human Connectome Project at humanconnectome.org, WU-Minn Consortium. The WU-Minn HCP Consortium obtained full informed consent from all participants, and research procedures and ethical guidelines were followed in accordance with the Washington University Institutional Review Boards (IRB #201204036; Title: ‘Mapping the Human Connectome: Structure, Function, and Heritability’). MATLAB (https://www.mathworks.com/) and SPSS (https://www.ibm.com/analytics/spss-statistics-software) were used to conduct the experiment’s reported in this study. The datasets supporting this article and the codes required to reproduce them can be found at https://github.com/longzhou-xu/data_and_code_sort.git.

## Results

### The signature of critical dynamics in the cortical network

For the 295 available subjects, we first investigated the power-law distribution of avalanche size at the population level. Here, we defined the activation as the time point when the BOLD signals reached their peak value, while the signals one step before and after this time point were above the chosen threshold (Fig. 1A). After preprocessing, the atlas_96_ signals were converted into point processes in which each time point represented an activation. We then calculated the avalanche size distribution *P*(*S*)~*S^−a^* (Fig. 1B), as well as the avalanche duration distribution *P*(*T*)~*T^−τ^* (Fig. 1C). First, from the estimated α and τ values, we tested whether the relationship between the scaling exponents holds for different thresholds of ROI signals (Fontenele et al., 2019; Friedman et al., 2012). We found that the closest matching occurred when the chosen threshold was around 1.4 SD (Fig. 1D). Second, the power-law distribution of avalanche sizes with a slope of *α* = 1.5 could be predicted by theory for a critical branching process with branching parameter *σ* = 1 (Harris, 1964; Zapperi, Lauritsen, & Stanley, 1995). However, we found that the threshold of 1.4 SD yielded σ = 0.91 and α = 1.56 (Fig. 1E), which did not match well with the theoretical prediction. We ran the same analysis on both atlas_246_ signals (Fig. S1a) and atlas_1024_ signals (Fig. S1b) to find the mismatch still exists (Fig. S1c-d).

As the above close check of hallmarks of criticality did not agree with each other well, we moved forward to investigate whether this mismatch could be a result of intersubject variability. We calculated both the MS and SE (see Synchrony and variability in synchrony) using the atlas_96_ signals for each of the 295 subjects and characterized the brain states of each subject with points in the MS vs. SE phase plane, as seen in the top panel of Fig. 2A. We found that the value of MS from these subjects extended from 0.2 to 0.7, and the distribution of subjects was not even but exhibited a greater tendency to the low MS range (Fig. 2A, bottom panel). This result suggested that even in the resting state, there is significant variability among the subjects’ brain states. It is clearly seen that these state points formed an inverted-U trajectory in the phase plane. The SE exhibited a maximum at the moderate value of MS, which implied the existence of a state with dynamic richness between order and disorder. We found the inverted-U curves and the calculation of SE were robust against different parcellation (Fig. S2). We also performed a phase randomization method on the fMRI data and found that this inverted-U curve disappeared in the randomized surrogate datasets (size=500, identified by visual inspection; examples can be seen in Fig. S3). Therefore, we argued that this inverted-U curve reflected a special spatiotemporal structure of brain dynamics that did not exist in randomized data.

**Figure 2.**
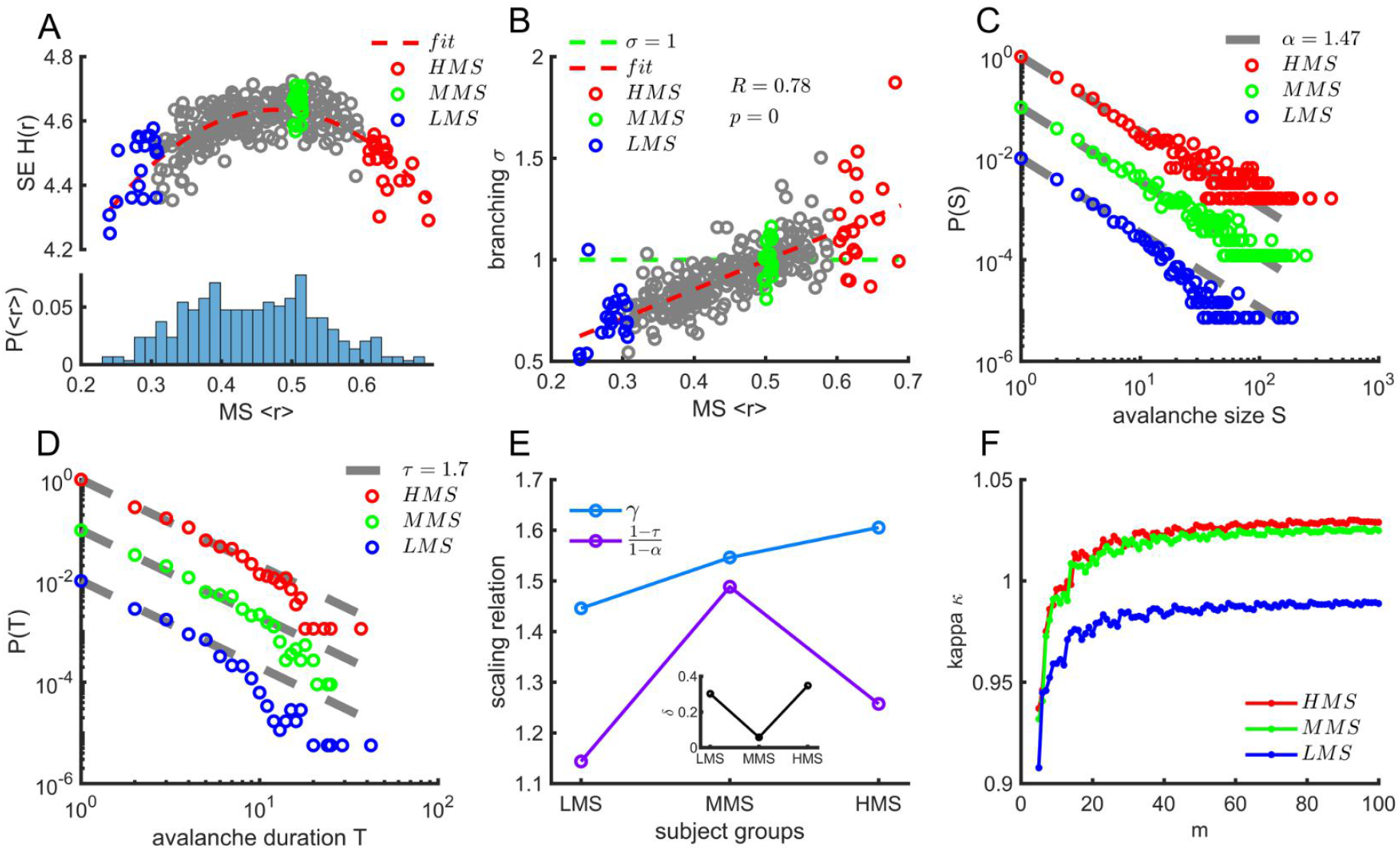
Signatures of criticality as a function of MS in resting-state brain networks. **A.** Top panel: The inverted-U trajectory of the MS 〈*r*〉 vs. SE *H*(*r*). The red dash line represents the quadratic fit of the data (*F*=188.758, *p*<0.001, adjusted *R*^2^=0.561). Bottom panel: The frequency count for the distribution of MS 〈*r*〉. **B.** The branching parameters *σ* vs. 〈*r*〉 for each subject. The green dashed line indicates *σ* = 1. The Pearson correlation value *R* and the *p* value are shown in the figure. The red dashed line represents the linear regression. For further analysis, we selected three representative groups of subjects according to their synchronization level: namely, LMS group (〈*r*〉 = 0.2824 ± 0.0219, blue open circles in A and B), MMS group (〈*r*〉 = 0.5041 ± 0.0042, green open circles in A and B) and HMS group (〈*r*〉 = 0.6304 ± 0.0246, red open circles in A and B). **C.** Avalanche size distributions for the LMS group, MMS group, and HMS group. To show the difference between these groups, we used gray lines with *α* = 1.47 to guide the eyes. The corresponding group-aggregated branching parameters are *σ_LMS_* = 0.7237 for the LMS group, *σ_MMS_* = 1.0123 for the MMS group, and *σ_HMS_* = 1.2023 for the HMS group. **D.** Avalanche duration distributions for three groups in **C**. To show the difference between these groups, we used gray lines with *τ* = 1.7 to guide the eyes. **E.** Scaling relations for the three groups. The blue line and purple line correspond to *γ* and 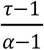, respectively. In the inset, 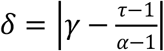 indicates the distance to the critical point. **F.** The dependence of *κ* on the numbers *m* of histogram bins for the three groups.

As expected, with the increasing of MS, the spatiotemporal activation pattern defined before exhibited transitions from random states to ordered states (Fig. S4). We then calculated the branching parameter *σ* for each subject. We found that with increasing MS, the branching parameter increased from less than 1 to higher than 1, crossing 1 at a moderate value of MS (Fig. 2B, and Fig. S5 for different parcellation).

Furthermore, we selected three groups from the above subjects: the low mean synchronization group (LMS group; the 20 most left subjects in Fig. 2A with an MS value of 〈*r*〉 = 0.2824 ± 0.0219), the moderate mean synchronization group (MMS group; the 20 subjects located near the peak of curve in Fig. 2A with an MS value of 〈*r*〉 = 0.5041 ± 0.0042), and the high mean synchronization group (HMS group; the 20 most right subjects in Fig. 2A with an MS value of 〈*r*〉 = 0.6304 ± 0.0246). For each group, we performed avalanche distribution analysis to identify which group was closest to the critical point (Fig. 2C-F). After obtaining scaling exponents *α* and *τ* for each group with a threshold of 1.4 SD (Fig. 2C-D), the scaling relationship showed the best match for the MMS group (Fig. 2E), and more detailed analysis results can be found in Fig. S6 in the Supplementary Materials.

Previous study showed that the truncations of power-law fit have a dramatic impact on power-law exponents, particularly on the ratio 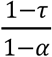, while *γ* barely changes (Destexhe & Touboul, 2020). To test the robustness of our results, we performed analysis in Fig 2c-e with different fitting windows for avalanche size S (*S_min_* ∈ [1, 10], and *S_max_* ∈ [30, 60]) and avalanche duration T (*T_min_* ∈ [1, 5], and *T_max_* ∈ [9, 20]). We found that the ratio of the number of fittings that met the critical criterion (|(1 − *τ*)/(1 − *α*) − *γ*| < 0.1 and Clauset’s goodness-of-fit test *p* > 0.1) to all power-law-fit samples is highest for MMS group (*φ* = 0.0346 for LMS, *φ* = 0.1103 for MMS, *φ* = 0.0266 for MMS).

We also calculated *κ*, an often-used parameter that could distinguish the difference between data and the theoretically suggested power-law distribution (Fagerholm et al., 2015; Palva et al., 2013; Poil, Hardstone, Mansvelder, & Linkenkaer-Hansen, 2012;Shew et al., 2009; Shew et al., 2011). As shown in Fig. 2F, as the discrete bin number *m* increases, the *κ* values become stable. The stabilized *κ* is smaller than 1 for the LMS group but larger than 1 for the HMS and MMS groups. The *κ* value for MMS group was closest to 1. Therefore, the above results suggested that subjects’ brains with moderate MS and maximal SE are poised closest to the critical point, supported by consistent hallmarks of criticality. On the other hand, the large dispersion of subjects among the phase space between asynchronous (subcritical) and synchronous (supercritical) states also provides an opportunity to investigate the phase transition in brains.

### The complexity in the FC network is maximized by criticality

Since the disorder-order phase transition could be observed, we investigated how this phase transition could impact the organization of FC networks. For convenience, we used MS to indicate this transition. We assessed how the variousness in FC strength changes as the brain undergoes a phase transition from the sub- to supercritical states. We used FC entropy and FC diversity as measures of variousness in FC strength in the brain networks. FC entropy is a direct measure of Shannon entropy from the probability distribution of FC strength obtained from the FC matrix, whereas FC diversity measures the similarity between the distribution of real FC matrix elements and uniform distribution. In previous studies, the former had been associated with healthy aging (Yao et al., 2013), and the latter is predicted to be maximized at the critical point by a computer model with Ginzburg-Landau equations (R. Wang et al., 2019). We found that both FC entropy (Fig. 3A) and FC diversity (Fig. 3B) peaked at the moderate value of MS; however, the peak position for these two measures was more rightward than that of SE.

**Figure 3.**
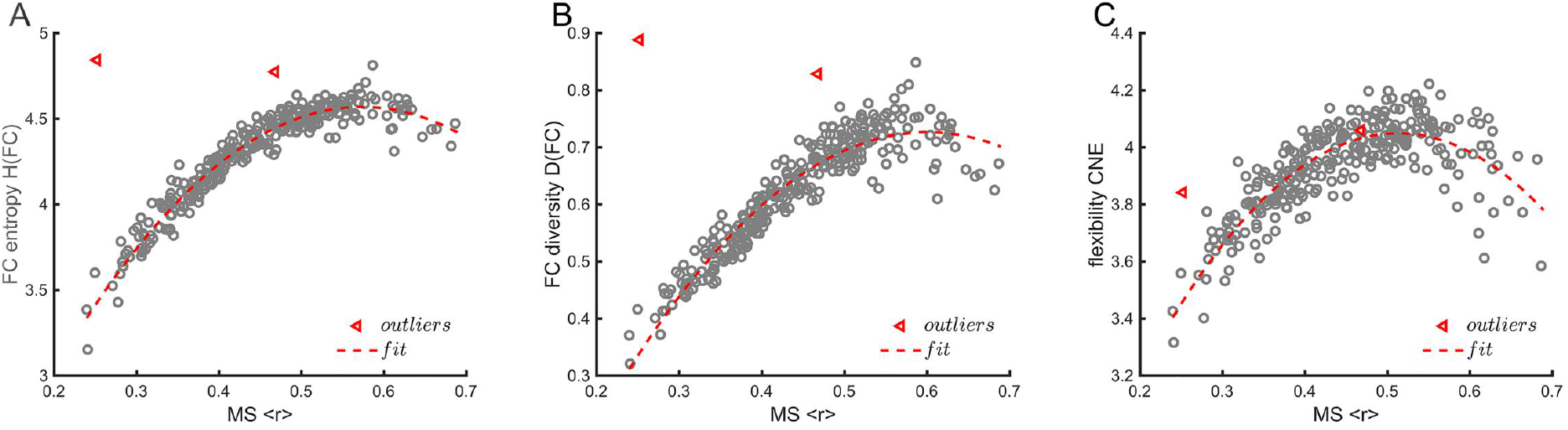
Dependence of complexity in the FC network on the MS of BOLD signals. **A.** FC entropy *H*(*FC*) as a function of MS 〈*r*〉. Red dashed line: quadratic fitting (*F* = 2287.892, *p*<0.001, adjusted *R*^2^ = 0.940). **B.** FC diversity *D*(*FC*) as a function of MS 〈*r*〉. Red dashed line: quadratic fitting (*F* = 1226.057, *p*<0.001, adjusted *R*^2^ = 0.894). **C.** FC flexibility *CNE* as a function of MS 〈*r*〉. Red dashed line: quadratic fitting (*F* = 366.851, *p<0.001*, adjusted *R*^2^ = 0.715). The red open triangles represent participants with outliers in the quadratic fittings in **A-B**.

The flexibility in dynamic FC reflects the extent of abundant connection patterns among regions and how frequent switching may occur between different patterns. In this work, we adopted the connection number entropy (*CNE*) as a measure of flexibility in FC networks. Our previous study showed that this measure was maximized at the critical point in a large-scale brain network model that combined DTI structural data and excitable cellular automaton (Song et al., 2019), and this measure could be reduced in the brains of patients with moyamoya disease (Lei et al., 2020). In this study, we found that the flexibility in FC was maximized with a moderate value of MS (Fig. 3C). The maximization was robust in a wide range of *THR_FC_* thresholds and sliding window lengths (Fig. S7). This result supported our previous conclusion (Lei et al., 2020; Song et al., 2019). Compared with FC entropy and FC diversity, the peak position for FC flexibility was nearer to the critical point.

FC entropy, diversity, and flexibility are often used in rfMRI studies to measure the complexity in the structure and dynamic reconfiguration of FC networks. Here, the study suggested that the complexity in FC networks is maximized by criticality.

### The maximized structure-function coupling around the critical point

From the obtained FC matrix and SC matrix for each subject, we constructed the FC networks for each subject and a group-aggregated SC network (see Similarity between functional and structural networks). We used *THR_FC_* and *THR_SC_* to control the link density in the FC networks and the group-aggregated SC network, respectively. We measured the similarity between the FC network and group-aggregated SC network with Pearson correlation and Hamming distance. Fig. 4A and B demonstrate the dependence of similarity on the MS of each subject with a link density of 0.7 in the FC networks. The similarity between the FC and SC was maximal for subjects with moderate synchrony, as the Pearson correlation was maximized (Fig. 4A), while the Hamming distance was minimized (Fig. 4B) for these subjects. This maximization of similarity between the FC and SC could be observed in a wide range of link densities in the FC and SC networks. To further consolidate the above results, we measured the similarity between the FC and SC for the three groups (LMS MMS and HMS) defined above as a function of the FC network link density. Fig. 4C-D show that as the FC network link density increased, the correlation coefficient between the FC and SC matrices first increased and then decreased, and consistently, the Hamming distance exhibited the opposite tendency. When the FC link density was large, the MMS group showed a significantly higher correlation and a lower Hamming distance between the FC and SC networks than the other two groups. Similarly, by varying *THR_SC_*, we found that the maximized similarity in the FC and SC at the critical point was robust in the wide range of link densities in the SC network (Figs. S8, S9 and S10).

**Figure 4.**
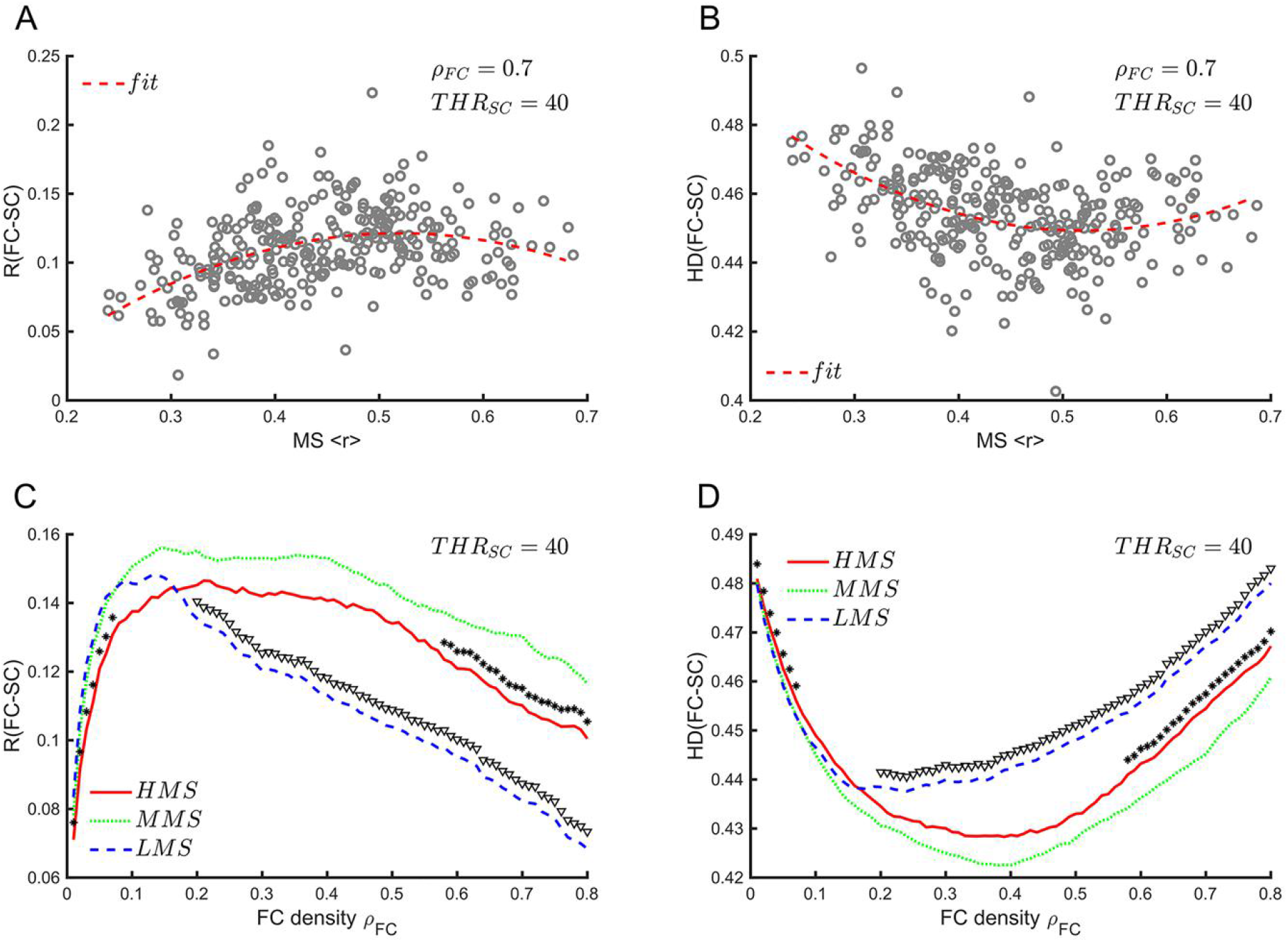
The dependence of structure-function coupling on the MS of brain networks. **A.** Pearson correlation between anatomical and functional networks as a function of MS 〈*r*〉. The link density in the FC network *ρ_FC_* = 0.7 and threshold in the group-aggregated SC network *THR_SC_*=40 (corresponding to *ρ_SC_* = 0.4836) are shown in the figure. Red dashed line: quadratic fitting (*F* = 36.997, *p* < 0.001, adjusted *R*^2^ = 0.197), which is better than linear fitting (*F* = 39.346, *p* < 0.001, adjusted *R*^2^ = 0.115). **B.** Hamming distance *HD*(*FC* − *SC*) between anatomical and functional networks as a function of MS 〈*r*〉. Red dashed line: quadratic fitting (*F* = 36.997, *p* < 0.001, adjusted *R*^2^=0.197), which is better than linear fitting (*F* = 39.346, *p* < 0.001, adjusted *R*^2^ = 0.115). **C.** The Pearson correlation between anatomical and functional networks as a function of FC density *ρ_FC_* (*THR_SC_* = 40 corresponding to *ρ_SC_* = 0.4826) for the HMS, MMS, and LMS groups. **D**. The Hamming distance between anatomical and functional networks as a function of FC density *ρ_FC_* (*THR_SC_* = 40 corresponding to *ρ_SC_* = 0.4826) for the HMS, MMS, and LMS groups. In **C-D**, the stars indicate significant differences between the HMS and MMS groups (two-tails two-sample t-test, *p* < 0.05, uncorrected); the open triangles indicate significant differences between the LMS and MMS groups (two-tails two-sample t-test, *p* < 0.05, uncorrected).

We noticed that for a large link density of the FC network, the dependence of similarity on link density monotonically decreased (Fig. 4C-D; Fig. S8a-b). Since lower link density conserved only stronger links in FC networks, we deduced that structural connections were mostly reflected in the strong functional connections. Meanwhile, the similarity also decreased as the threshold *THR_SC_* in SC networks decreased (Fig. S9a-b), suggesting that the structural connections that were mostly reflected in the functional connections were those shared by most subjects because structural connections specified to individuals would be excluded with high *THR_SC_*.

### The dynamic phase transition in individual subjects’ brains

The observed individual brain states dispersed around the critical point provided an opportunity to investigate the dynamic phase transition in individual brains. To this end, for the LMS, MMS, and HMS groups defined above, we randomly selected two subjects from each group. We calculated the dynamical MS 〈r〉_n_ and SE H(r)_n_ for these six subjects with the sliding window approach (Fig. 5A, s1-s6) from their Kuramoto order parameters r(t) (Fig. 5B). We observed a time-dependent change in individuals’brain states in the state space following the inverted-U trajectory, as shown in the top panel of Fig. 2A. In the time period limited by scan duration, we observed that subjects who were farther away from the critical point tended to stay in the regime decided by MS, and events of crossing the critical point (black lines at r(t) = 0.5) to the other regime seldom occurred (s1, s2, s5, and s6 in Fig. 5A, or Fig. 5B, top and bottom panel). Subjects who were nearer the critical point were more likely to cross the critical point (s3 and s4 in Fig. 5A, or Fig. 5B, middle panel).

**Figure 5.**
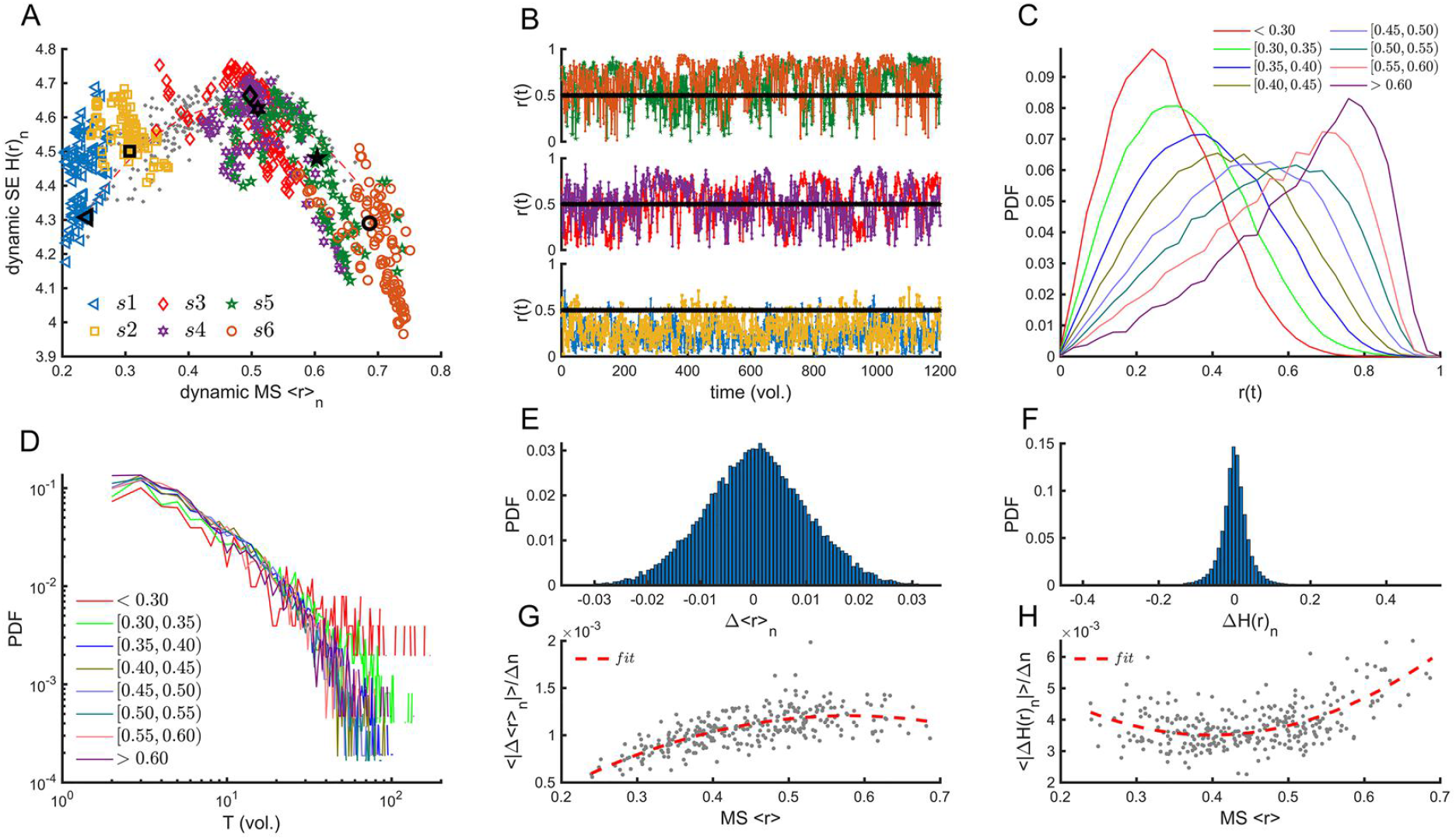
The dynamic phase transition in individual subjects’ brains. **A.** The dependence of dynamic SE *H*(*r*)_*n*_ on dynamic MS 〈*r*〉_*n*_ from six subjects selected randomly from the LMS, MMS, and HMS groups. The enlarged dark markers indicate the mean position for corresponding subjects (markers with the same shape). **B.** The time-dependent changes in the Kuramoto order parameter *r*(*t*) for six subjects as demonstrated in figure a (with the same color). **C.** The normalized frequency count of *r*(*t*) for different levels of 〈*r*〉, indicated by lines with different colors. **D.** The dwell time (the time interval between two successive critical point crossing events) distribution for different levels of 〈*r*〉. **(E, F).** The distribution of vertical and horizontal moving distances of phase points in one step of the sliding window. **(G, H).** The vertical and horizontal velocities of state points of each subject as a function of their MS 〈*r*〉. The vertical and horizontal velocities were calculated by 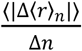 and 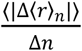, where the symbol |·| indicates the absolute value, and 〈·〉 was the average across all the windows. Δ*n* is the step used to slide the windows. Here, Δn=10 time points (volumes). Red dashed lines in **G-H**: quadratic fitting (*F* = 139.316, *p* < 0.001, adjusted *R*^2^ = 0.485 in (G); *F* = 81.181, *p* < 0.001, adjusted *R*^2^ = 0.353 in figure h). Both quadratic fittings were better than linear fitting (adjusted *R*^2^ = 0.407 in **G** and adjusted *R*^2^ = 0.173 in **H**).

To validate the above observation at the population level, we divided the 295 subjects at hand into eight groups with different levels of synchrony and calculated the corresponding probability distribution of the Kuramoto order parameter *r*(*t*). It is seen clearly from Fig. 5C that as the synchrony level decreases, the distribution of the Kuramoto order parameter becomes narrow and less tilted. Meanwhile, we found that the dwell time, which referred to the time interval between two successive critical point crossing events, exhibited heavier tails in its distribution for low synchrony groups (Fig. 5D). These results implied the higher inertness in the subcritical regime than others, and brains were more likely to stay in this regime with longer dwell times.

Next, we calculated the distribution of vertical and horizontal moving distances in state space in a fixed time interval Δ*n* (the time points or volumes of one step of the sliding window) for all subjects. We found that the distributions of vertical and horizontal moving distances were both symmetrical with a mean of zero (Fig. 5E-F), suggesting that the inverted-U trajectory in the state space was stable and unlikely to change its shape as time progressed. Furthermore, the position (〈*r*〉) dependent velocity distribution is maximal for horizontal velocity 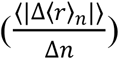 and minimal for vertical velocity 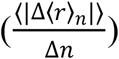 near the critical point (Fig. 5G-H). The maximal horizontal velocity around the critical point implied that at this point, the systems were most sensitive to the perturbations due to internal fluctuations or external modulations. Meanwhile, the lower vertical and horizontal velocities in the subcritical regime compared to the supercritical regime also reflected the high inertness in the subcritical regime.

It was of interest to determine whether the maximization of FC complexity, as well as function-structure coupling, around the critical point could be realized dynamically when the individual brains endured phase transition. To this end, we obtained the time-dependent FC matrices with the sliding window method and calculated FC entropy (Fig. 6A), FC diversity (Fig. 6B), and two measures for similarity between FC and SC (Fig. 6C-D) as a function of instantaneous MS in each time window. The time-dependent complexity and similarity measures followed almost the exact trajectories as those in the static measurements shown in Fig. 3A-B, as well as Fig. 4A-B. This result implied that FC complexity and similarity between FC and SC were indeed modulated by phase transition in brains, and their maximization could be realized dynamically by positioning the system around the critical point.

**Figure 6.**
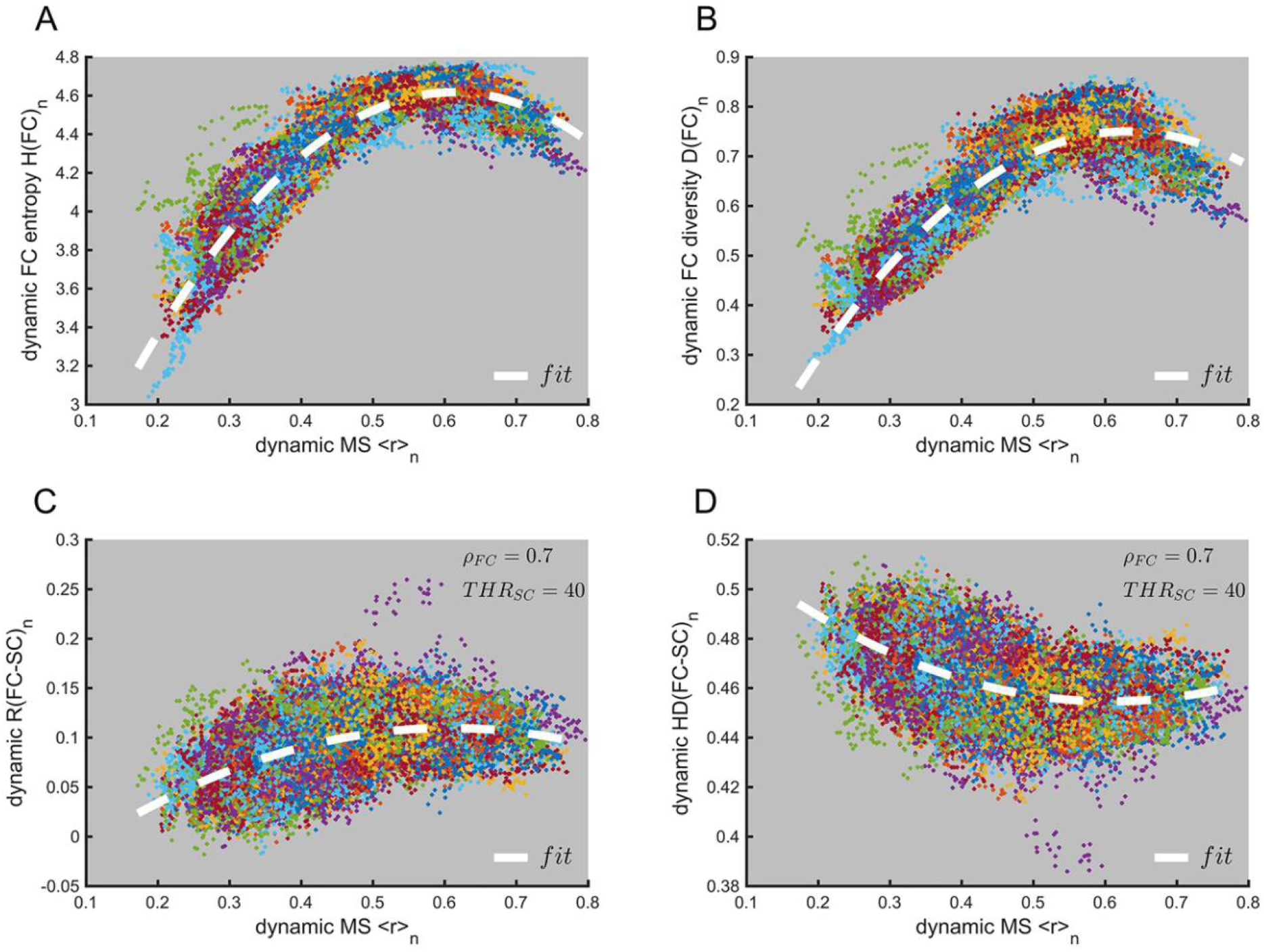
Dynamic modulations of FC complexity and structure-function coupling during the phase transition of brains. **A.** The dependence of dynamic FC entropy as a function of instantaneous MS; thick dashed white line: quadratic fitting (*F* = 106350.82, *p* < 0.001, adjusted *R*^2^ = 0.877). **B.** The dependence of dynamic FC diversity as a function of instantaneous MS. Thick dashed white line: quadratic fitting (*F* = 80261.492, *p* < 0.001, adjusted R^2^ = 0.843). **C.** The dependence of dynamic FC-SC correlation as a function of instantaneous MS; thick dashed white line: quadratic fitting (*F* = 5519.072, *p* < 0.001, adjusted *R*^2^ = 0.270), which is better than linear fitting (adjusted *R*^2^ = 0.225). **D.** The dependence of dynamic FC-SC Hamming distance as a function of instantaneous MS; thick dashed white line: quadratic fitting (*F* = 5519.072, *p* < 0.001, adjusted *R*^2^ = 0.270), which is better than linear fitting (adjusted *R*^2^ = 0.225). In **A-D**, each dot represents a calculation from one window. The dots with the same color represent the calculation for one subject. However, due to the limited number of colors used, different subjects may share the same color. In **C-D**, a link density of 0.7 was used to obtain the binary FC network, and a threshold of 40 was used to obtain the group-aggregated structural network.

### High fluid intelligence and working memory capacity were associated with critical dynamics

The results above support the hypothesis that large-scale brain networks lie in the vicinity of a critical point which is associated with moderate MS and maximal SE. Another key prediction from the critical brain hypothesis is that brains that are closer to criticality should be better in cognitive performance. Here, to address this prediction, we assessed linear relationships between SE and intelligence scores of the available subjects. We found that SE values were significantly correlated with fluid intelligence scores (PMAT; Fig. 7A) but not with crystallized intelligence scores (picture vocabulary; Fig. 7B). Meanwhile, we found that working memory scores, which were assessed using the Listing Sorting Working Memory test from the NIH Toolbox, were significantly correlated with SE (List sorting; Fig. 7C). We also noted here that these scores were significantly correlated with many other measures that were found to be maximized at the criticality, namely, FC entropy, FC diversity, and FC flexibility (Fig. S11 in Supplementary Materials). Meanwhile, there were significant quadratic relationships between MS and fluid intelligence, as well as working memory scores, but not for crystallized intelligence scores (Fig. 7D-F). We also found these results still held when potential confounds such as age and education achievements were regressed out (Fig. S12 and S13). Therefore, these results imply that brains that are closer to criticality are associated with higher fluid intelligence and working memory scores.

**Figure 7.**
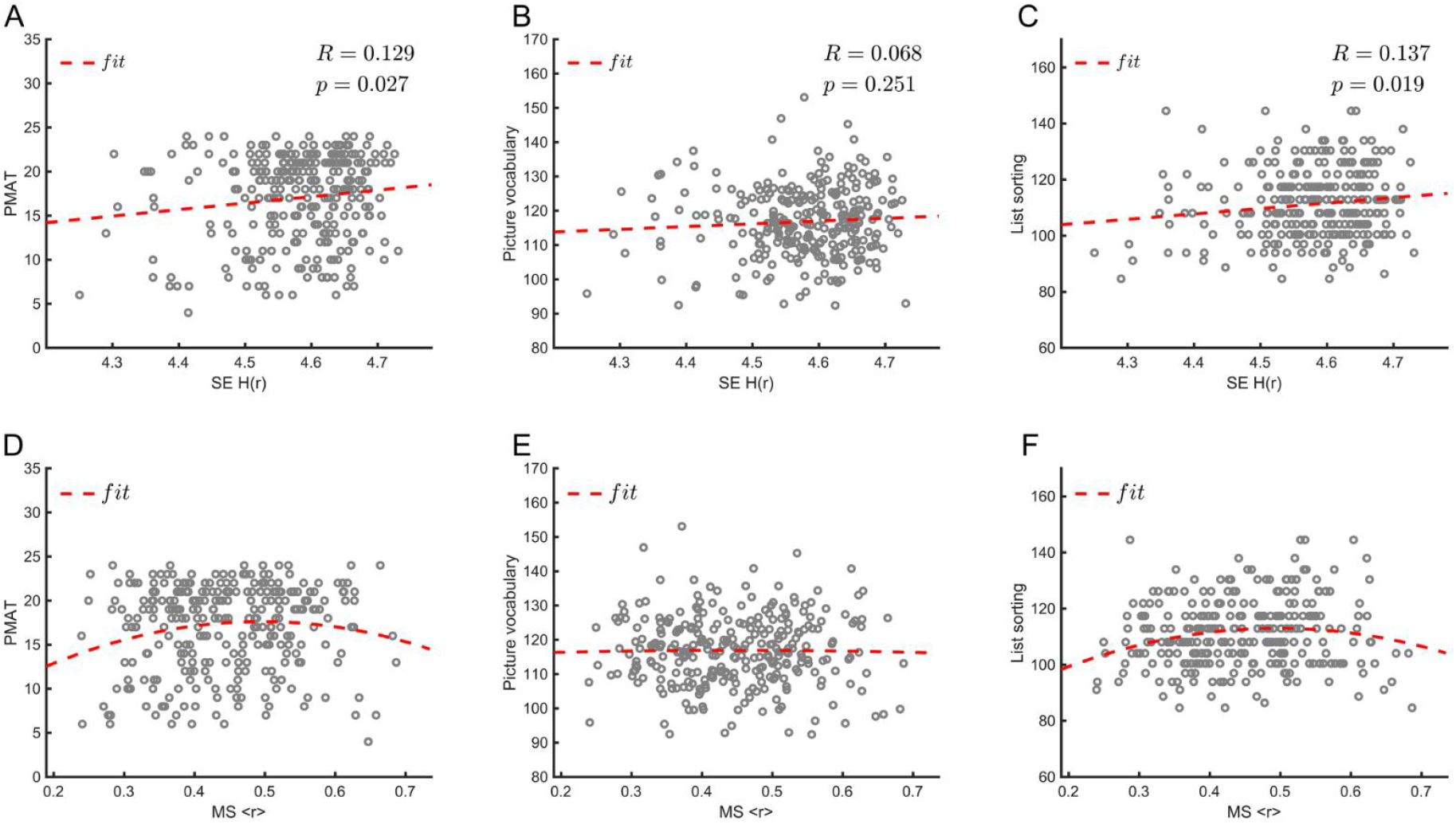
Correlations between cognitive performance scores and SE, as well as MS. **A-C.** Correlation between SE and PMAT scores, picture vocabulary test scores, as well as the list sorting working memory test scores. Red dashed lines in **A-C**: linear fitting. **D.** Scatterplot of the PMAT scores against the MS. The red dashed line represents the significant quadratic fit of the data (*F* = 3.145, *p* = 0.045, adjusted *R*^2^ = 0.015), which is better than the linear fitting (adjusted *R^2^* = 0.004). **E.** Scatterplot of the picture vocabulary test scores against the MS. Both the linear and quadratic regressions are not significant (linear: *p* = 0.991; quadratic: *p* = 0.988). **F.** Scatterplot of the list sorting working memory test scores against the MS. The red dashed line represents the significant quadratic fit of the data (*F* = 4.376, *p* = 0.013, adjusted *R*^2^ = 0.023), which is better than linear fitting (adjusted *R*^2^ = 0.008).

Since a wide variety of experiments have demonstrated that fluid intelligence is associated with a distributed network of regions in the Parieto-Frontal Integration Theory (P-FIT), including frontal areas (Brodmann areas (BAs) 6, 9, 10, 45-47), parietal areas (BA 7, 39, 40), visual cortex (BAs 18, 19), fusiform gyrus (BA 37), Wernicke’s area (BA 22) and dorsal anterior cingulate cortex (BA 32) (Jung & Haier, 2007; Nikolaidis et al., 2017), we decided to find more relationships between these regions with critical dynamics indicated by maximized SE.

To obtain the relevant regions in a fine-grained division of the brain, here we used the Human Brainnetome Atlas, which contains 210 cortical and 36 subcortical subregions (Fan et al., 2016). We extracted from each brain region the voxel-level BOLD signals and calculated the regional SE for these 246 regions. We found that regions whose SE exhibited significant (*p* < 0.05, FDR corrected) positive correlations with PMAT scores were located in the frontal areas (i.e., bilateral SFG, MFG, PrG, right IFG and PCL), parietal areas (i.e., bilateral AG, SMG, Pcun, right SPL), right inferior temporal gyrus (ITG), superior occipital gyrus (sOcG) and left Cingulate gyrus (CG) (Fig 8 and Table 1).

**Figure 8.**
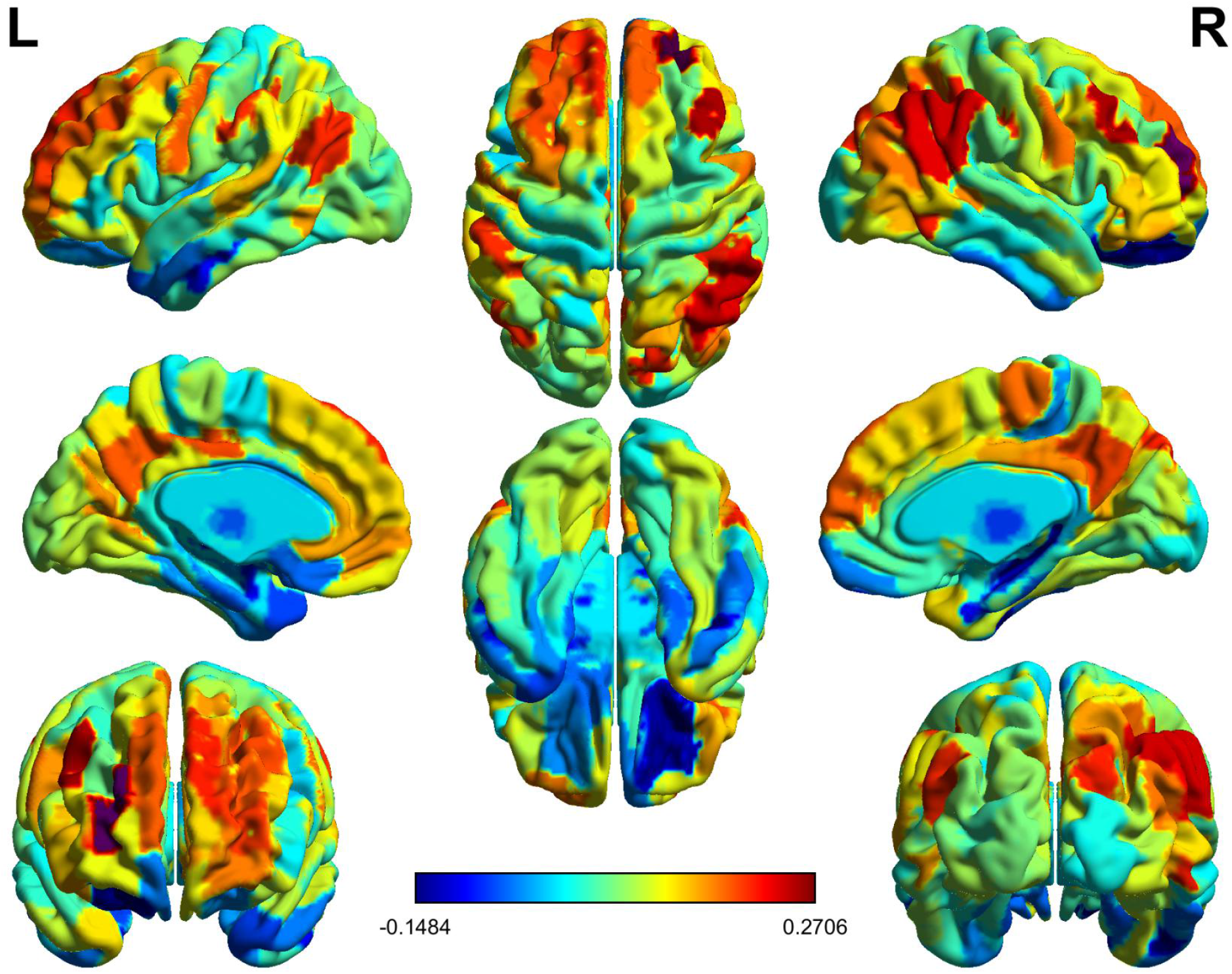
The brain map for correlations between regional SE and fluid intelligence. The color bar indicated the Pearson correlation value between regional SE and PMAT. The cortical and subcortical regions were defined by the Human Brainnetome Atlas (http://atlas.brainnetome.org/bnatlas.html). Data was visualized using BrainNet Viewer (Xia, Wang, & He, 2013).

**Table 1.**
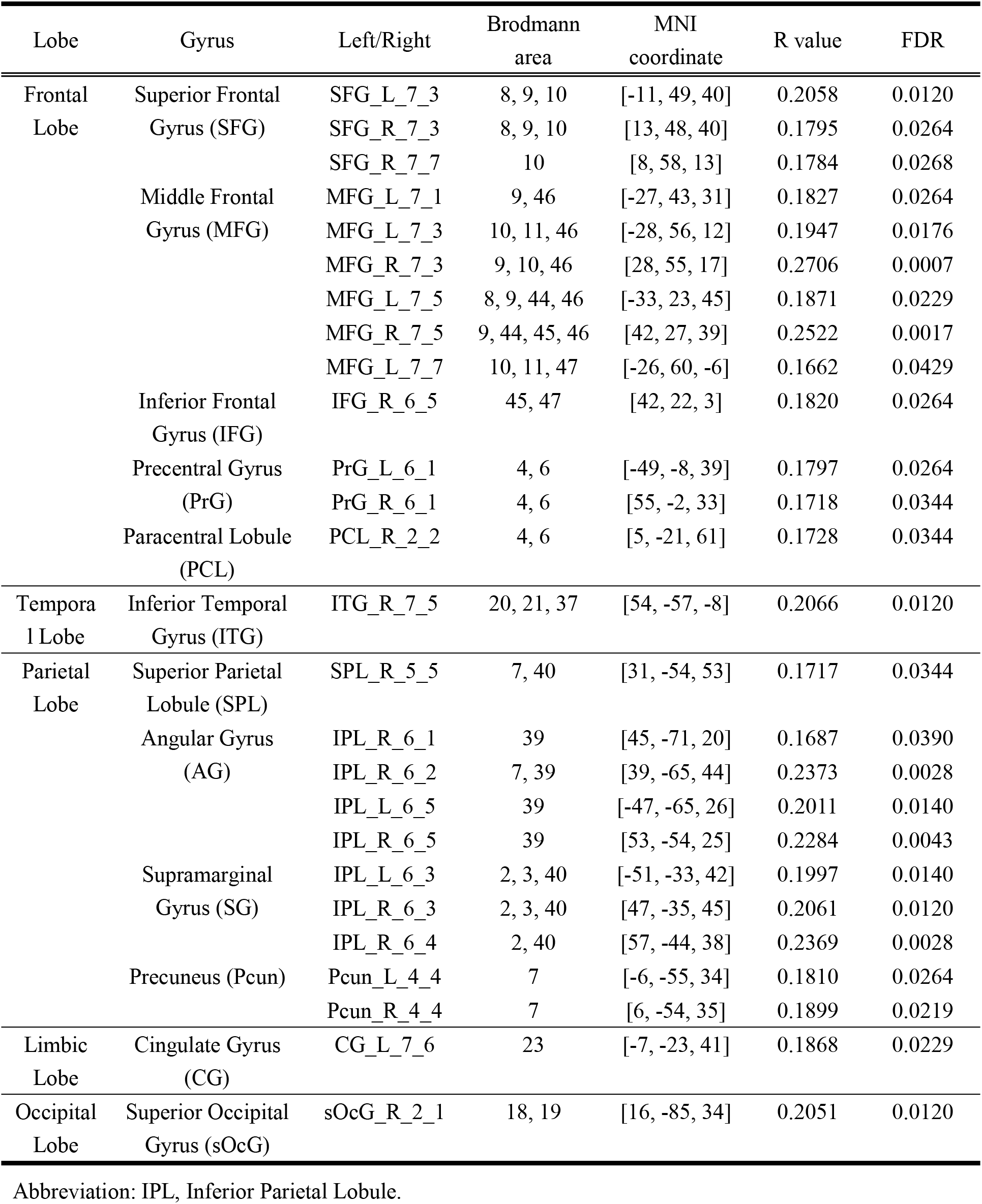
The brain regions exhibited significant correlation between SE and fluid intelligence.

## Discussion

In this study, using the large scale WU-Minn HCP dataset and a large number of criticality-inspired metrics, we provided evidence that though healthy young brains at rest are on average close to the critical point, as critical brain hypothesis has suggested, there is a considerable individual variation around this point. This gave us a chance to validate previous theoretical predictions of criticality on large scale brain networks, and indeed we observed that the complexity of brain FC, as well as the structure-function coupling, was maximized around the critical point. We proceeded to observe a dynamic phase transition in individual subjects, and found that their brains tended to stay subcritical, as indicated by a longer dwell time in this parameter region. Finally, we found that high fluid intelligence and working memory capacity were associated with critical dynamics rather than noncritical dynamics, not only globally but also regionally, suggesting the functional advantages of critical dynamics in resting-state brains.

Balance between functional segregation and functional integration is a central organizing principle of the cerebral cortex. It has been argued that FC complexity characterizes the interplay of functional segregation and functional integration (Sporns, 2013). A comparison between simulated and empirically obtained resting-state FC indicates that the human brain at rest lies in a dynamic state that reflects the largest complexity its anatomical connectome can host (G. Tononi, Sporns, & Edelman, 1994). Recently, many studies have tried to link complexity with cognitive performance, human intelligence, and even consciousness, either measured by Φ (big phi) in integrated information theory or discriminated between levels of sedation (Ahmadlou et al., 2014; Duncan, Chylinski, Mitchell, & Bhandari, 2017; Saxe, Calderone, & Morales, 2018; Giulio Tononi, Edelman, & Sporns, 1998). Meanwhile, there is a growing awareness that complexity is strongly related to criticality. A recent study showed that criticality maximized complexity in dissociated hippocampal cultures produced from rats (Timme et al., 2016). Here, in this study, we measured FC complexity from different perspectives, either on its strength diversity or on its dynamic flexibility (Fig. 3A-C, Fig. 6A-B). With the observation of the phase transition trajectory, we demonstrated that these measures of FC complexity were maximized around the critical point. Therefore, the formulation that criticality maximizes complexity was supported in our work empirically with fMRI data at the whole-brain network level.

It has been shown that human brains possess a stable set of functionally coupled networks that echo many known features of anatomical organization (Krienen, Yeo, & Buckner, 2014). Several computational modeling studies have demonstrated that critical dynamics could best explore the repertoire provided by the structural connectome (Deco & Jirsa, 2012; Enzo Tagliazucchi et al., 2016). Recent studies also suggested the capacity of repertoire provided by the structural connectome could be extended by the hierarchical modular structural organization (R. Wang et al., 2019). Therefore, structure-function coupling was believed to be at its maximal when the system is at criticality (R. Wang et al., 2019), and it could be disrupted by losing criticality (Cocchi et al., 2014), or disruption of hierarchical organization of structural networks. Previous studies in anesthetized human brains have found structure-function decoupling accompanied by unidirectional departure from a critical point (Enzo Tagliazucchi et al., 2016). It is possible that functional connectivity flexibility could be used as a measure of the extent that functional connectivity explores the repertoire provided by structural connectome, and the highest functional connectivity flexibility occurs when the system is at criticality (Song et al., 2019) (Fig. 3C). Our work demonstrated the maximal exploration of structural connections at the critical point occurs in resting state brains (Fig. 4). However, since we used a group aggregated structural connection networks, we did not investigate how organization of structural connections could impact on the capacity of network repertoire. This issue will be investigated in the future.

Interestingly, although the brain hovers around the critical point, the brain prefers to stay in the subcritical region, as the subject distribution was skewed toward a disordered state, and the dwell time in the subcritical state was longer (Fig. 5). Previous analysis of in vivo data has argued that the mammalian brain self-organizes to a slightly subcritical regime (Priesemann et al., 2014). It was suggested that operating in a slightly subcritical regime may prevent the brain from tipping over to supercriticality, which has been linked to epilepsy.

Meanwhile, with a slightly subcritical regime deviates only little from criticality, the computational capabilities may still be close to optimal. However, our results showed that the resting state brains could actually stay in the supercritical regimes. So, the preference of brains for subcritical regime may not because of prevention of too ordered states. In another study, by relating the EEG-domain cascades to spatial BOLD patterns in simultaneously recorded fMRI data, the researchers found that while resting-state cascades were associated with an approximate power-law form, the task state was associated with subcritical dynamics (Fagerholm et al., 2015). They argued that while a high dynamic range and a large repertoire of brain states may be advantageous for the resting state with near-critical dynamics, a lower dynamic range may reduce elements of interference affecting task performance in a focused cognitive task with subcritical dynamics (Fagerholm et al., 2015). Therefore, there remains a possibility that the resting state is not “pure resting state”, but mixed with some occasional “task state” for some subjects. However, further delicately designed experimental studies are required to test this conjecture. It remains to uncover the relationship between cognitive states and neural dynamics that lies on a spectrum. The method proposed in this study may be useful in future studies of this topic.

Recently, Ezaki et al. used the Ising model to map BOLD signals on a two-dimensional phase space and found that human fMRI data were in the paramagnetic phase and were close to the boundary with the spin-glass phase but not to the boundary with the ferromagnetic phase (Ezaki, Fonseca dos Reis, Watanabe, Sakaki, & Masuda, 2020). Since the spin-glass phase usually yields chaotic dynamics whereas the ferromagnetic phase is nonchaotic, their results suggested that the brain is around the “edge of chaos criticality” instead of “avalanche criticality”. However, our findings support that avalanche criticality occur in large-scale brain networks. Therefore, it is interesting to investigate whether both kinds of criticality could co-occur in large-scale brain networks (Kanders et al., 2017). Ezaki et al. also found that criticality of brain dynamics was associated with human fluid intelligence, though they used performance IQ to reflect fluid intelligence, which refers to active or effortful problem solving and maintenance of information. In our work, we assessed the correlation between fluid intelligence and the critical dynamics indicated by synchronization entropy for brain regions, and found regions showed significant positive correlations were located in parietal-frontal network (Fig. 8 and Table 1). These regions were most frequently reported in studies of intelligence and its biological basis, including structural neuroimaging studies using voxel-based morphometry, magnetic resonance spectroscopy, and DTI, as well as functional imaging studies using positron emission tomography (PET) or fMRI (Jung & Haier, 2007). Also, in the Parieto-Frontal Integration Theory of intelligence, these regions are considered as the most crucial nodes of the brain network underlying human intelligence (Jung & Haier, 2007; Nikolaidis et al., 2017).

Our study suggested that not only fluid intelligence, but also working memory capacity was associated with critical dynamics. This is possibly because working memory may share the same capacity constraint through similar neural networks with fluid intelligence (Halford, Cowan, & Andrews, 2007; Jaeggi, Buschkuehl, Jonides, & Perrig, 2008; Kane & Engle, 2002). In our study, the critical dynamics in the frontal and parietal network also exhibited significant correlation with working memory capacity (Fig. S14 and Table S1). Furthermore, it has been well established that working memory is strongly modulated by dopamine, and too strong or too weak dopamine D1 activation is detrimental for working memory, with the optimal performance achieved at an intermediate level (Cools & D’Esposito, 2011; Vijayraghavan, Wang, Birnbaum, Williams, & Arnsten, 2007; Zahrt, Taylor, Mathew, & Arnsten, 1997). This inverted-U dose-response has been observed in mice (Lidow, Koh, & Arnsten, 2003), rats (Zahrt et al., 1997), monkeys (Cai & Arnsten, 1997) and humans (Gibbs & D’Esposito, 2005). Recent studies on neural network models have shown that the optimal performance of working memory co-occurs with critical dynamics at the network level and the excitation-inhibition balance at the level of individual neurons and is modulated by dopamine at the synaptic level through a series of U or inverted-U profiles (Hu, Huang, Jiang, & Yu, 2019). Here in this study, we demonstrated that the optimal performance of working memory and criticality co-occurs at the system level.

However, our study had several limitations. Firstly, the surrogate data test used in this study ruled out the possibility that the results we obtained could be explained by autocorrelations in the data. However, the long-range spatial correlation of criticality cannot allow one to test the results by ruling out the effects of correlation across the time series. Secondly, though we used the denoising fMRI data from HCP with standard data pre-processing procedure, it is still interesting to investigate how the pre-processing procedure affects the results. Thirdly, in the avalanche analysis, the activation events defined in this study were slightly different from definition used by others, such as threshold-crossing events (Enzo Tagliazucchi et al., 2012) or above-threshold events (Bocaccio et al., 2019; R. Wang et al., 2019). We compared these different methods and found all these methods could generate scale free avalanche activities, but unlike our method, the other two methods failed to generate critical branching process (See Section II in Supplementary Materials). Therefore, it is interesting to investigate the correlations between neural activities and events detected by different detection methods from BOLD signals. Finally, in this study we only focused on the cognitive abilities that are associated with critical dynamics, and found significant but not strong correlations between fluid intelligence, working memory and critical dynamics. Recent works demonstrated that the functional network segregation, integration and their balance could predict different cognitive abilities (R. Wang et al.,2021). Therefore, future investigation on the relationship between this functional balance and criticality across individuals may reveal various associations between diverse cognitive abilities with not only critical dynamics, but also non-critical dynamics. Despite all these shortages, we hope this study may inspire future research work to validate our findings, e.g., through observing not only the association between the departure of criticality and the decline of cognitive performance, either in aging or brain disease, but also the restore of criticality and the improvement of cognitive performance with pharmacological or noninvasive brain stimulation (Barch, 2004; Reinhart & Nguyen, 2019).

## Conclusions

In conclusion, we mapped individuals’ brain dynamics from resting-state fMRI scans on the phase transition trajectory and identify subjects who are close to the critical point. With this approach, we validated two predictions of critical brain hypothesis on large-scale brain networks, i.e., maximized FC complexity and maximized structure-function coupling around the critical point. We also observed the tendency of brain to stay in subcritical regime. Finally, we found that the critical dynamics in large-scale brain networks were associated with high scores in fluid intelligence and working memory, implying the vital role of large-scale critical dynamics in cognitive performance. We also identified key brain regions whose critical dynamics was highly correlated with human intelligence. Our findings support the critical brain hypothesis that neural computation is optimized by critical brain dynamics, as characterized by scale-free avalanche activity, and could provide a solution for improving the effects of future interventions targeting aspects of cognitive decline (Reinhart & Nguyen, 2019), possibly by controlling the criticality through non-invasive stimulation (Chialvo, Cannas, Grigera, Martin, & Plenz, 2020).

