## Supplementary material for "Avalanche criticality in individuals, fluid intelligence and working memory": supplymental figures and table

**for**

### Section I: Supplementary Figures

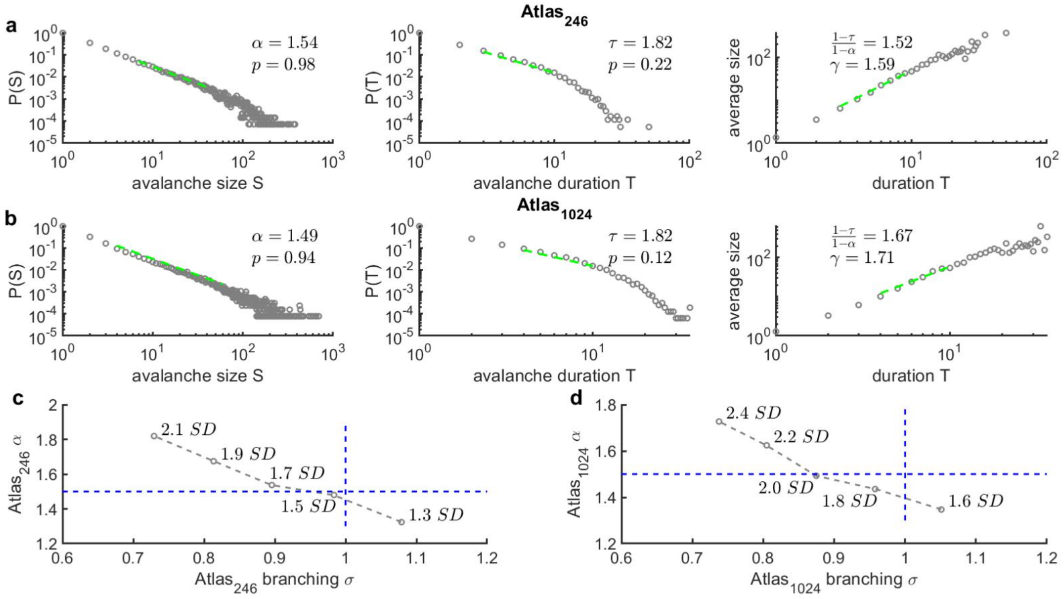


**Figure S1. Avalanche statistics obtained from group-level analysis using atlas with different size.** The power-law distributions of avalanche sizes (left panel), avalanche durations (middle panel), and the power-law relationship between avalanche sizes and durations (right panel) for atlas$_{246}$ (a), atlas$_{1024}$ (b). Note that here the events were defined as the suprathreshold peak intermediate between two above-threshold time points (threshold = 1.7 SD for atlas$_{246}$, 2 SD for atlas$_{1024}$), which were same as Figure 1. The power-law exponents of avalanche sizes α and avalanche durations $\tau$ are also shown, as well as the Cluset's test results $p$. The power-law exponent $\gamma$ of the relationship between avalanche sizes and durations is predictable from a theoretical formula $\frac{1-\tau}{1-\alpha}$ at different resolutions.

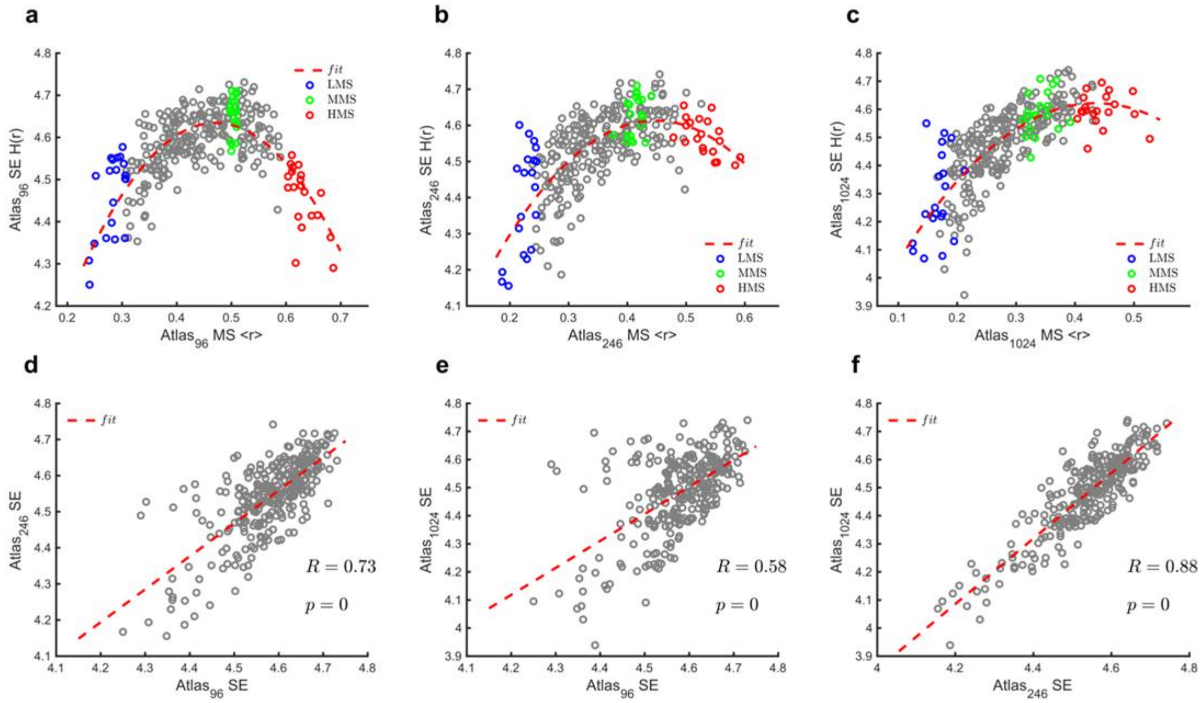


**Figure S2. The robustness of inverted-U curve between MS and SE for atlas with different size. a-c.** The global MS and SE calculated from extracted ROI BOLD signals using atlas$_{96}$ (**a,** $F = 188.758$, $p$<0.001, adjusted $R^2 = 0.561$), atlas$_{246}$ (**b,** $F = 145.994$, $p$<0.001, adjusted $R^2 = 0.497$) and atlas$_{1024}$ (**c,** $F = 217.496$, $p$<0.001, adjusted $R^2 = 0.596$). The red dashed lines are the quadratic regression. **d-f.** The SE calculated from different atlas was correlated, suggesting the subjects near the critical point could be successfully identified with different atlas. The Pearson correlation value $R$ and $p$ were shown on the figures. The red dashed lines are the linear regression.

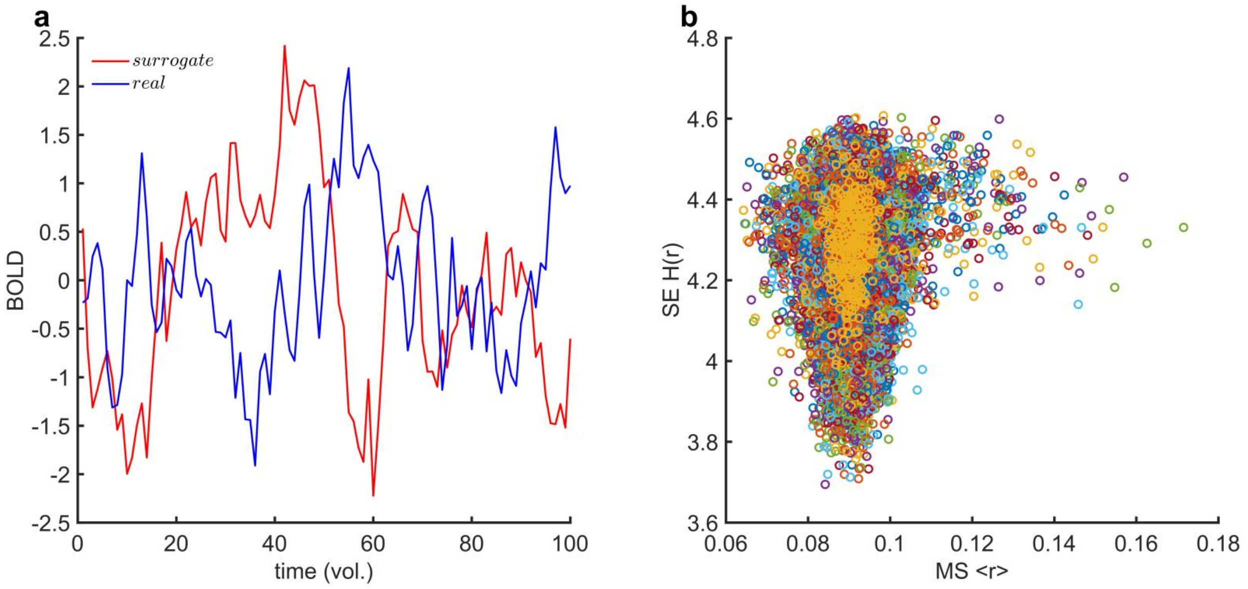


**Figure S3. To verify the calculated synchrony and entropy are not due to noise. a.** The plot contains examples of real ROI BOLD signals and surrogate signals from phase shuffling. **b.** The relationship between $\langle r \rangle$ and $\mathrm{H}(r)$ obtained from the phase shuffling. Note that we generated 500 phase-shuffling surrogate time series to obtain $\langle r \rangle$ and $\mathrm{H}(r)$. Different colors represent different surrogate data in this figure.

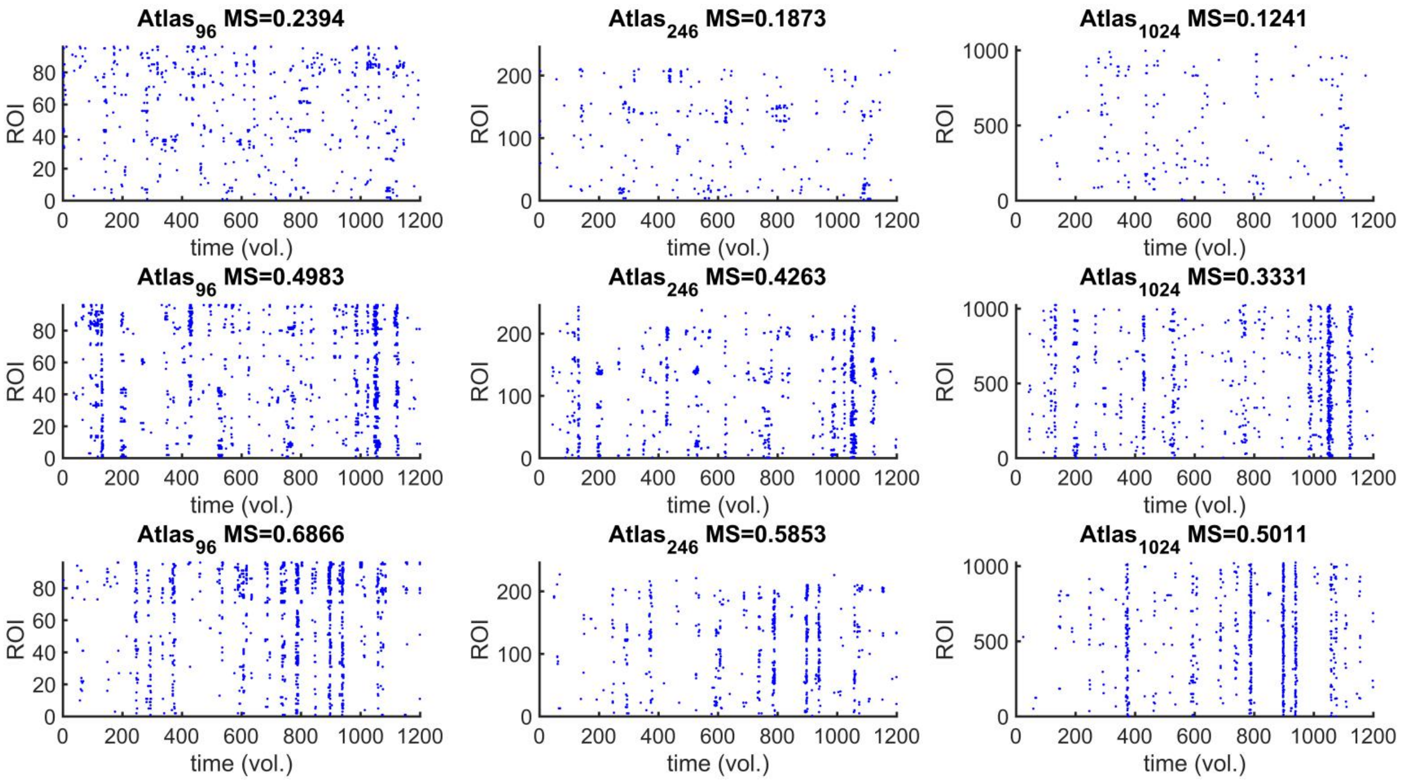


**Figure S4. Sample of event activity across the three levels of MS.** Top/Middle/Bottom panel: The event raster for the same participant with low/middle/high MS in three brain atlases. Note that here the events were defined as the suprathreshold peak intermediate between two above-threshold time points (threshold = 1.4SD for $atlas_{96}$, 1.7SD for $atlas_{246}$, 2SD for $atlas_{1024}$), which were same as Figure 1.

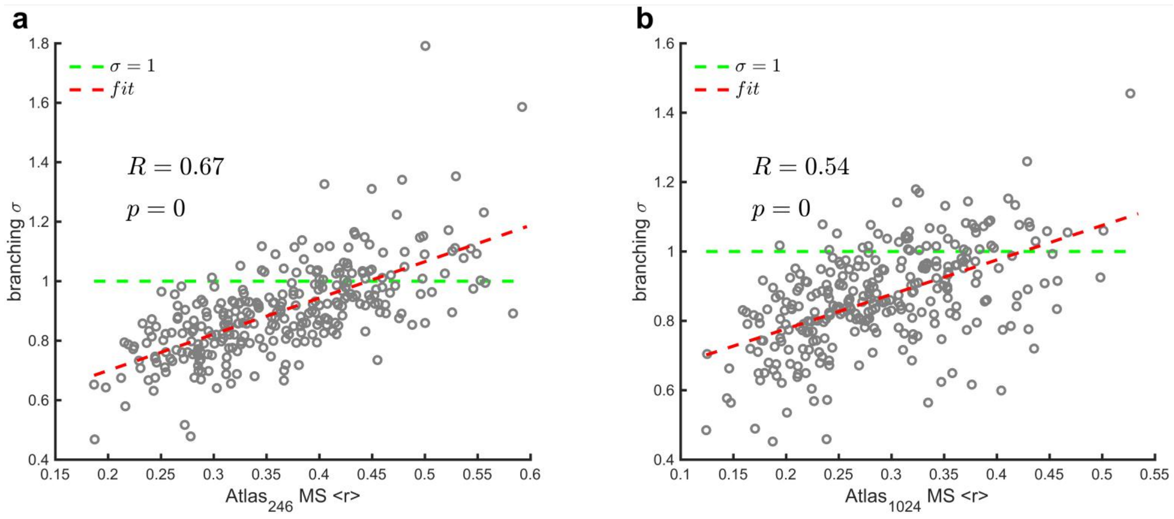


**Figure S5. The relationship between branching parameters and MS.** The branching parameters $\sigma$ vs. MS $< r >$ of each subject for atlas$_{246}$ (a) and atlas$_{1024}$ (b). The green dashed line indicates $\sigma = 1$. The Pearson correlation value $R$ and $p$ value are shown in the figure. The red dashed line is the linear regression. Note that here the events were defined as the suprathreshold peak intermediate between two above-threshold time points (threshold = 1.4SD for atlas$_{96}$, 1.7SD for atlas$_{246}$, 2.6SD for atlas$_{1024}$), which were same as Figure 1.

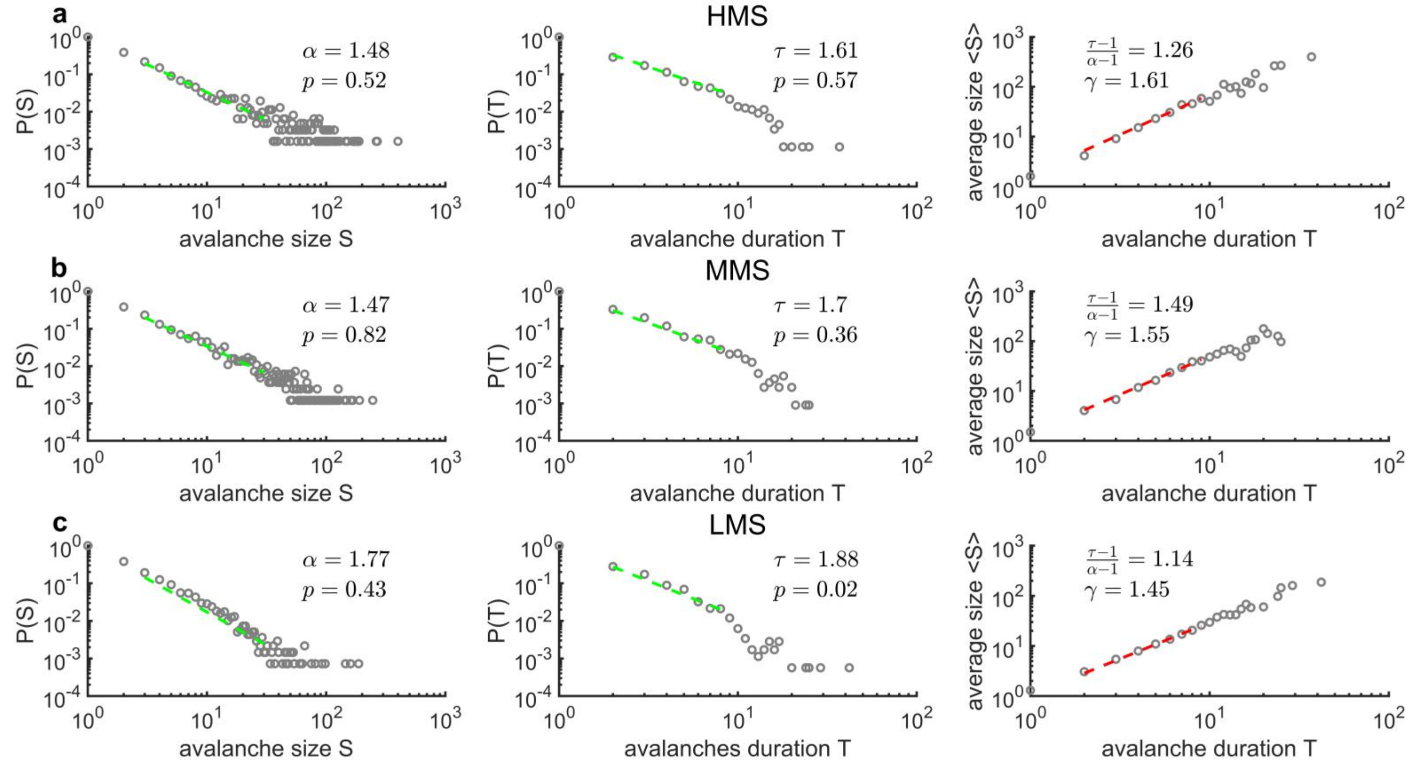


**Figure S6. Power-law distribution reflecting the brain dynamics for different levels of MS <r>.** The power-law distributions of avalanche size (left panel), avalanche durations (middle panel), and the power-law relationship between avalanche sizes and durations (right panel) in the HMS, MMS and LMS groups. Note that the truncations of power-law fit are same between three groups. The power-law exponents of avalanche sizes ($\alpha$) and avalanche durations ($\tau$) are also shown, as well as Clauset's test results ($p$). The power-law exponent ($\gamma$) of the relationship between avalanche sizes and durations is predictable from a theoretical formula $\frac{\tau-1}{\alpha-1}$ in the MMS group, reflecting MS closer to brain criticality.

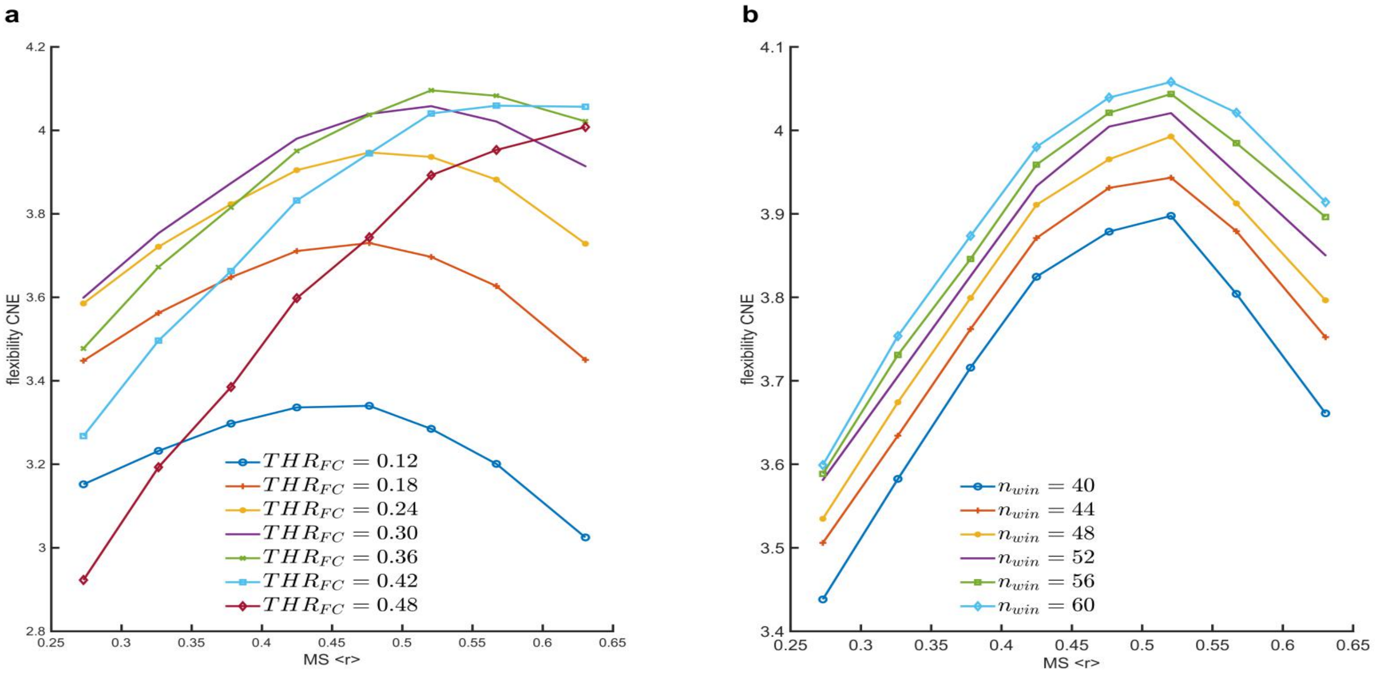


**Figure S7. The robustness of the association between flexibility and MS.** CNE was calculated using different thresholds for the FC matrix and different window lengths, and the data demonstrate that CNE is robust to some extent. **a.** The relationship between flexibility and MS $\langle r \rangle$ against different FC thresholds ($THR_{FC}$). **b.** The relationship between flexibility and MS against different numbers of windows ($n_{win}$).

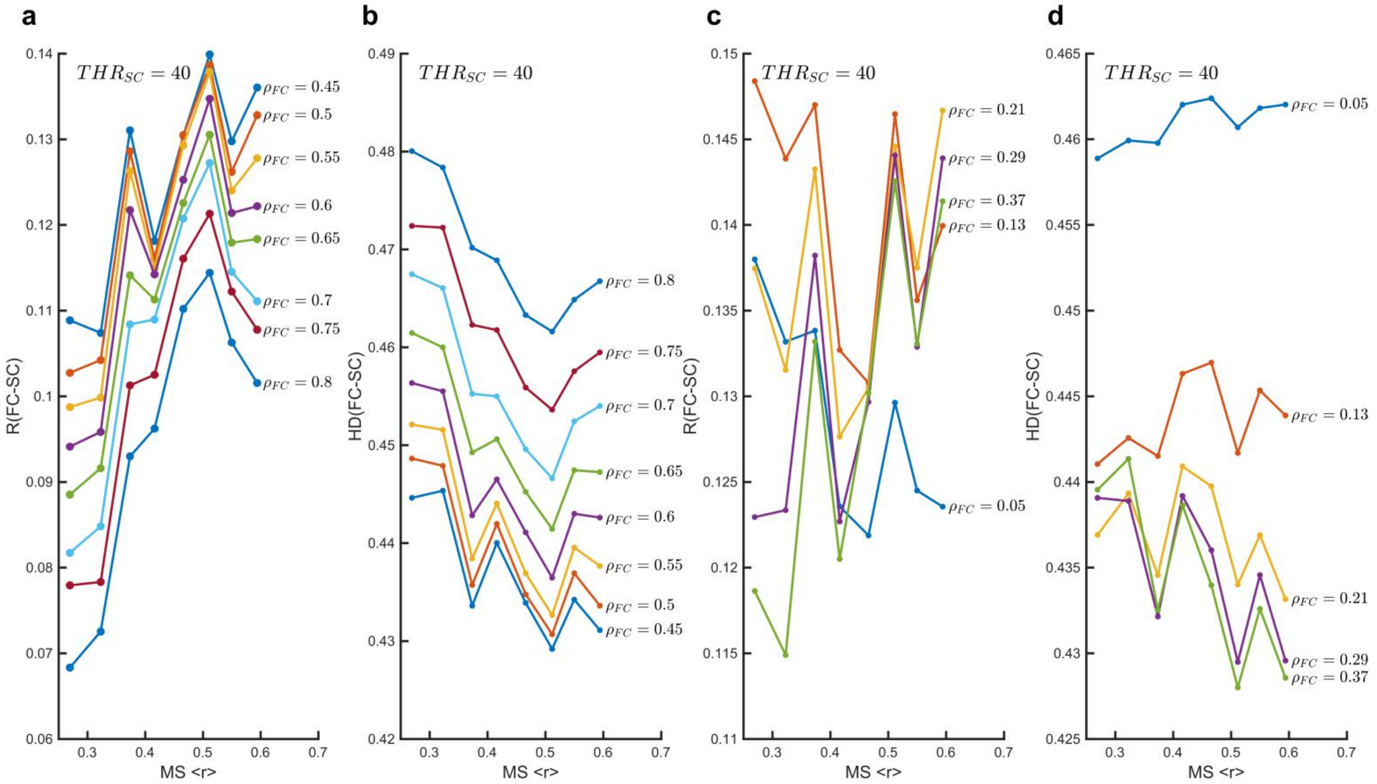


**Figure S8. The robustness of the similarity between FC and SC against different FC link densities. a and c.** Pearson correlations between SC and FC when the SC density is equal to 0.4837 ($THR_{SC} = 40$) against different FC link densities $\rho_{FC}$ (among 0.05-0.8). **b and d.** Hamming distance between SC and FC. Note that the solid circles indicate the average values.

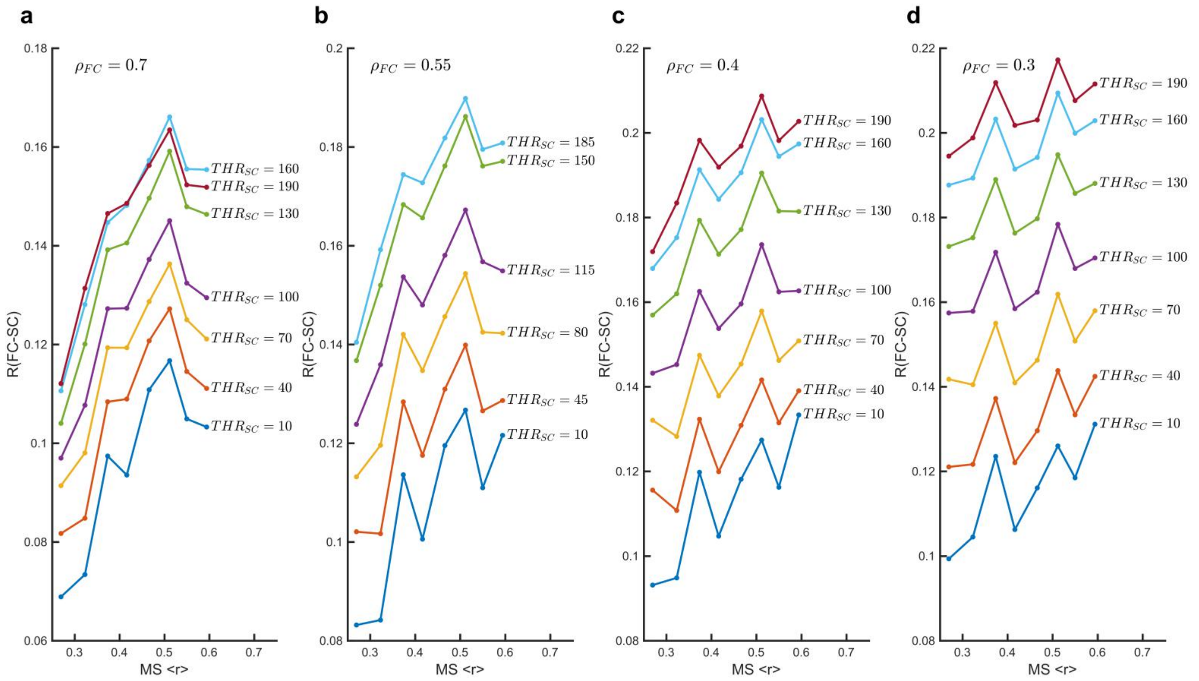


**Figure S9. The robustness of the Pearson correlations between FC and SC against different $THR_{SC}$. a.** Pearson correlation between SC and FC while the FC link density is equal to 0.7, against different $THR_{SC}$ (from 10 to 190, step is 30). **b.** Pearson correlations between SC and FC when the FC link density is equal to 0.55 against different $THR_{SC}$ (from 10 to 185, step is 35). **d.** Pearson correlations between SC and FC when the FC link density is equal to 0.4 against different $THR_{SC}$ (from 10 to 190, step is 30). **d.** Pearson correlations between SC and FC when the FC link density is equal to 0.3 against different $THR_{SC}$ (from 10 to 190, step is 30). Note that the solid circles indicate the average values.

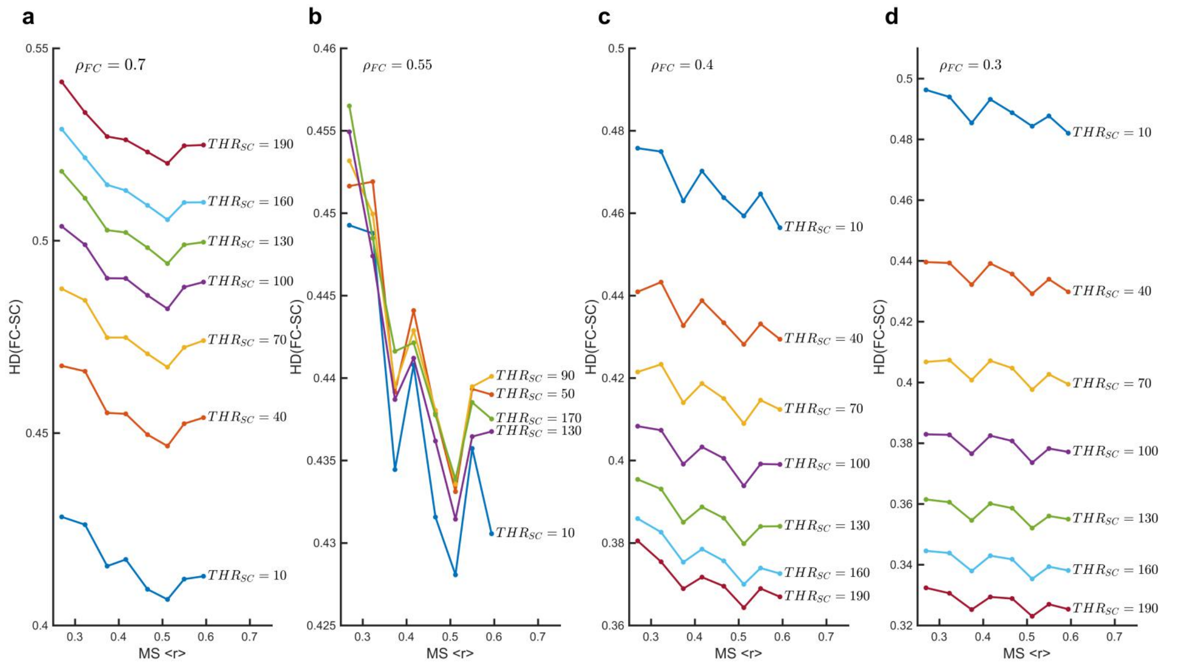


**Figure S10. The robustness of the Hamming distance (HD) between FC and SC against different $THR_{SC}$. a.** HD between SC and FC while the FC link density is equal to 0.7, against different $THR_{SC}$ (from 10 to 190, step is 30). **b.** HD between SC and FC while the FC link density is equal to 0.55, against different $THR_{SC}$ (from 10 to 170, step is 40). **c.** HD between SC and FC while the FC link density is equal to 0.4, against different $THR_{SC}$ (from 10 to 190, step is 30). **d.** HD between SC and FC while the FC link density is equal to 0.3, against different $THR_{SC}$ (from 10 to 190, step is 30). Note that the solid circles indicate the average values.

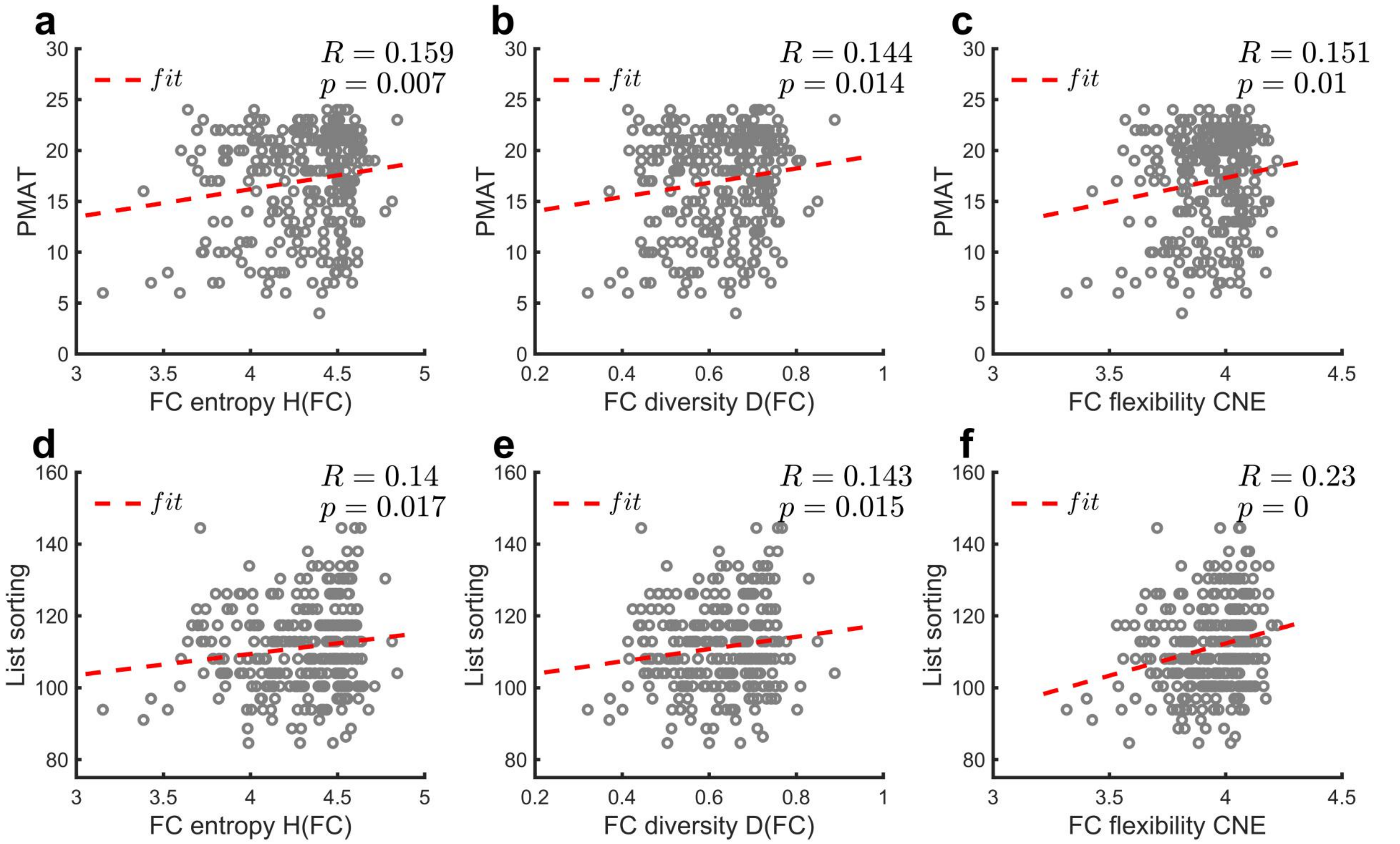


**Figure S11. Relationships between cognitive abilities and different FC complexity measures.** The Pearson correlation values (i.e., $R$) and $p$ values are shown in the figure. The red dashed lines represent the linear regression.

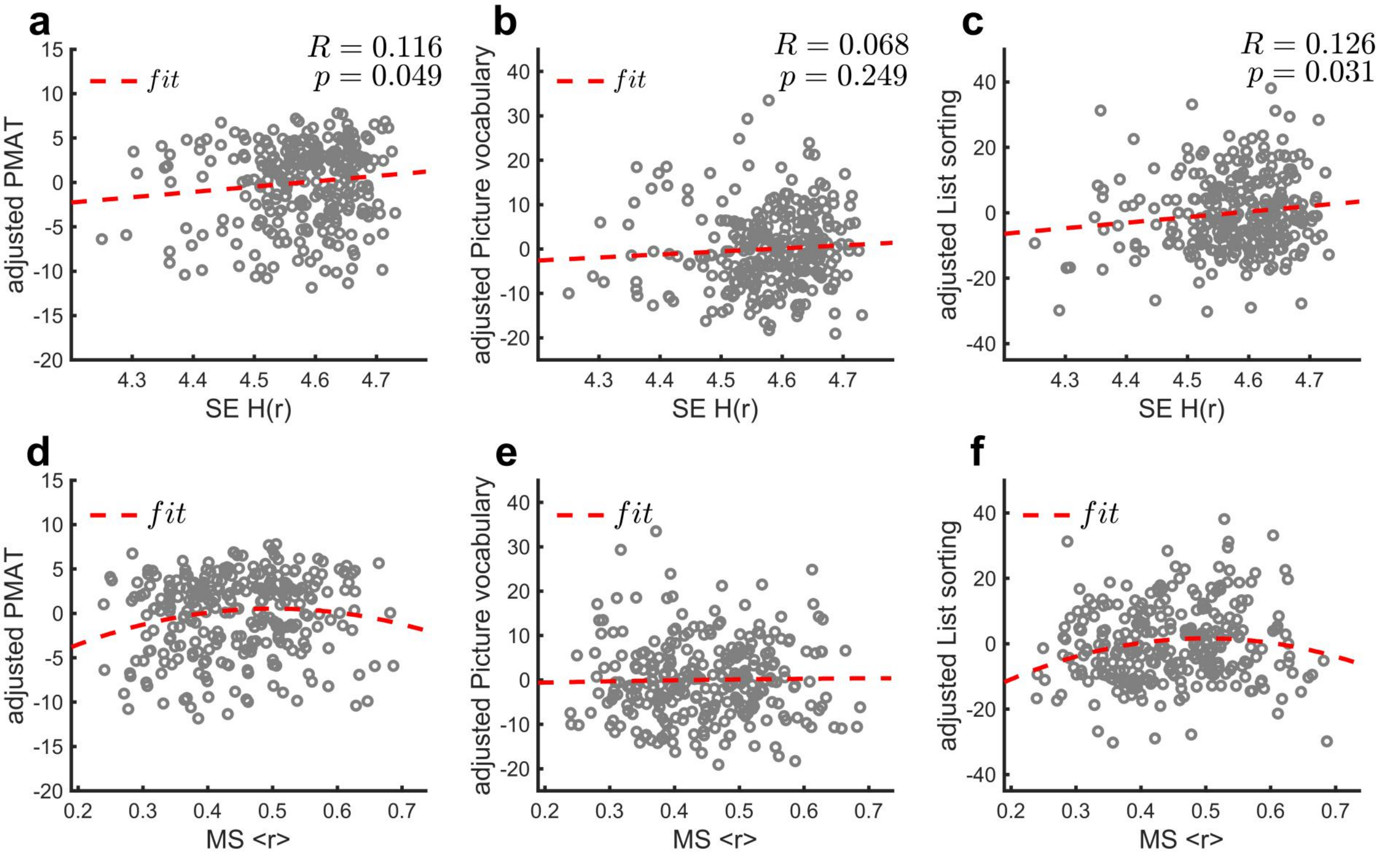


**Figure S12. Relationships between adjusted cognitive abilities and SE, as well as MS.** The adjusted cognitive scores were obtained by regressing out the age and education achievement from raw cognitive scores. **a-c.** Correlation between SE and adjusted PMAT scores, adjusted picture vocabulary test scores, as well as the adjusted list sorting working memory test scores. Red dashed lines in **a-c**: linear fitting. The Pearson correlation values (i.e., $R$) and $p$ values are shown in the figure. **d.** Scatterplot of the adjusted PMAT scores against the MS. The red dashed line represents the marginal significant quadratic fit of the data ($F = 2.871$, $p = 0.058$, adjusted $R^2 = 0.013$), which is better than the linear fitting (adjusted $R^2 = 0.005$). **e.** Scatterplot of the adjusted picture vocabulary test scores against the MS. Both the linear and quadratic regressions are not significant (linear: $p = 0.726$; quadratic: $p = 0.94$). **f.** Scatterplot of the adjusted list sorting working memory test scores against the MS. The red dashed line represents the significant quadratic fit of the data ($F = 4.095$, $p = 0.018$, adjusted $R^2 = 0.021$), which is better than linear fitting (adjusted $R^2 = 0.008$).

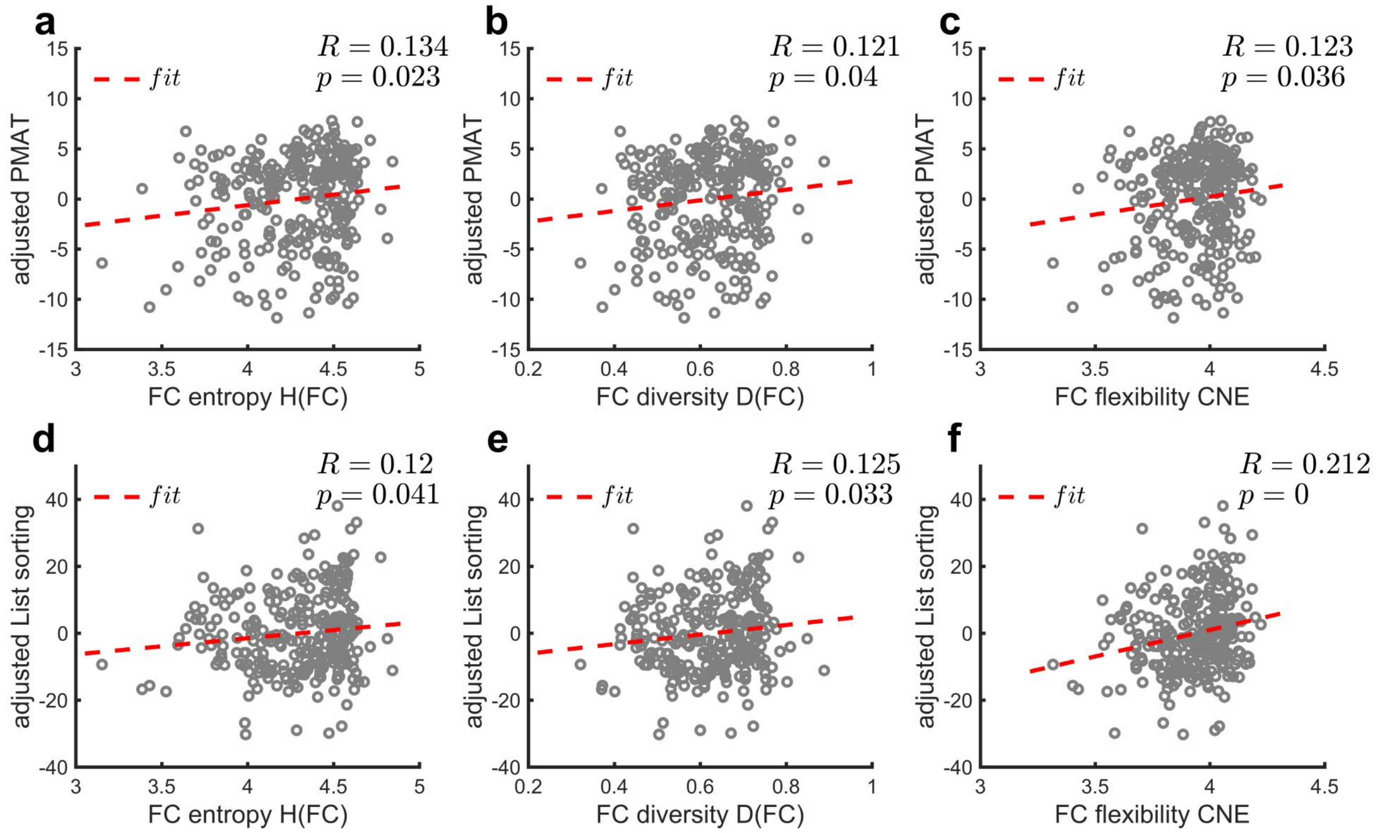


**Figure S13. Relationships between adjusted cognitive abilities and different FC complexity measures.** The adjusted cognitive scores were obtained by regressing out the age and education achievement from raw cognitive scores.

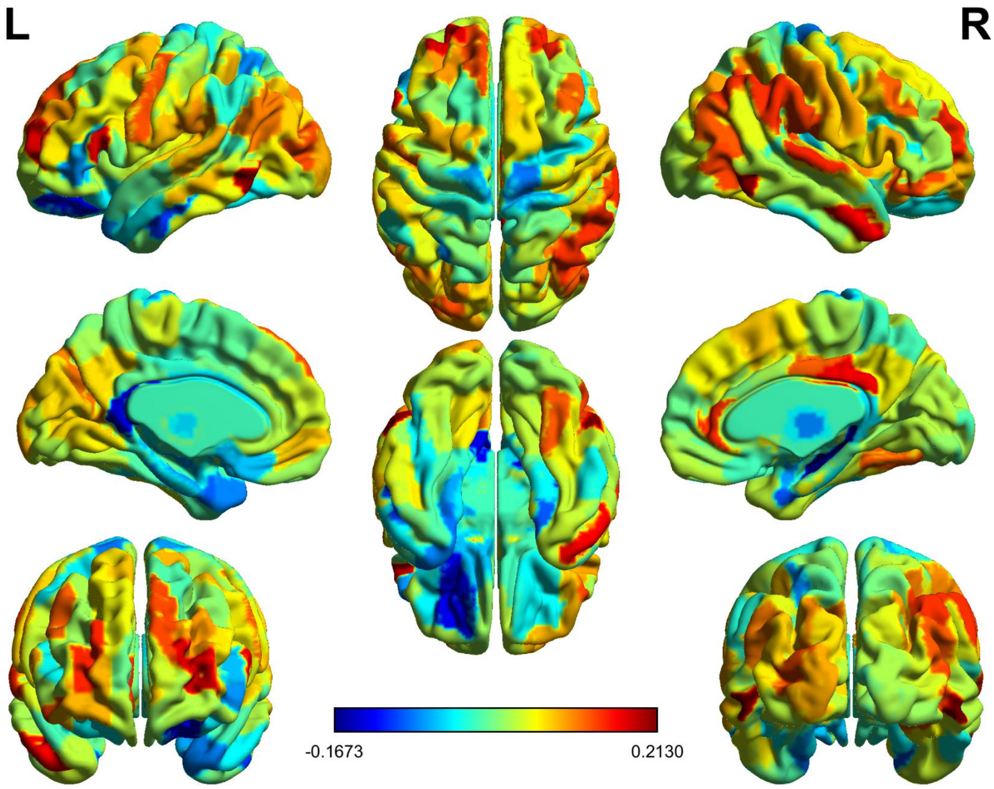


**Figure S14. The brain map for correlations between regional SE and List sorting work memory.** The color bar indicated the Pearson correlation value between regional SE and List sorting work memory. The cortical and subcortical regions were defined by the Human Brainnetome Atlas. Data was visualized using BrainNet Viewer.

**Table S1. The brain regions exhibited significant (uncorrected) correlation between SE and List**

**sorting work memory.**

| Lobe | Gyrus | Left/Right | MNI coordinate | R value | P value (uncorrected) |
|---|---|---|---|---|---|
| Frontal Lobe | Superior Frontal Gyrus (SFG) | SFG_L_7_3 | [-11, 49, 40] | 0.1411 | 0.0162 |
| | Middle Frontal Gyrus (MFG) | MFG_L_7_3 | [-28, 56, 12] | 0.1776 | 0.0024 |
| | | MFG_R_7_3 | [28, 55, 17] | 0.1546 | 0.0084 |
| | | MFG_R_7_5 | [42, 27, 39] | 0.1248 | 0.0336 |
| | Inferior Frontal Gyrus (IFG) | IFG_L_6_3 | [-53, 23, 11] | 0.1687 | 0.0040 |
| | Orbital Gyrus (OrG) | OrG_R_6_6 | [42, 31, -9] | 0.1444 | 0.0138 |
| | Precentral Gyrus (PrG) | PrG_L_6_1 | [-49, -8, 39] | 0.1399 | 0.0171 |
| Temporal Lobe | Superior Temporal Gyrus (STG) | STG_R_6_4 | [66, -20, 6] | 0.1530 | 0.0091 |
| | Middle Temporal Gyrus (MTG) | MTG_R_4_2 | [51, 6, -32] | 0.1642 | 0.0051 |
| | Inferior Temporal Gyrus (ITG) | ITG_L_7_5 | [-55, -60, -6] | 0.2070 | 0.0004 |
| | | ITG_R_7_5 | [54, -57, -8] | 0.2155 | 0.0002 |
| | Fusiform Gyrus (FuG) | FuG_R_3_2 | [31, -62, -14] | 0.1343 | 0.0222 |
| Parietal Lobe | Angular Gyrus (AG) | IPL_R_6_1 | [45, -71, 20] | 0.1350 | 0.0214 |
| | | IPL_R_6_2 | [39, -65, 44] | 0.1527 | 0.0092 |
| | | IPL_L_6_5 | [-47, -65, 26] | 0.1196 | 0.0418 |
| | Supramarginal Gyrus (SG) | IPL_R_6_4 | [57, -44, 38] | 0.1457 | 0.0130 |
| | | IPL_R_6_6 | [55, -26, 26] | 0.1360 | 0.0205 |
| | Precuneus (Pcun) | Pcun_L_4_3 | [-12, -67, 25] | 0.1254 | 0.0327 |
| Limbic Lobe | Cingulate Gyrus (CG) | CG_R_7_1 | [4, -37, 32] | 0.1649 | 0.0049 |
| | | CG_R_7_6 | [6, -20, 40] | 0.1316 | 0.0250 |
| | | CG_R_7_7 | [5, 41, 6] | 0.1500 | 0.0106 |
| Occipital Lobe | Occipital Gyrus (OcG) | OcG_L_4_1 | [-31, -89, 11] | 0.1325 | 0.0240 |
| | | OcG_R_4_2 | [48, -70, -1] | 0.1385 | 0.0183 |
| Subcortical Nuclei | Striatum (Str) | Str_L_6_3 | [-17, 3, -9] | 0.1190 | 0.0429 |

Abbreviations: IPL, Inferior Parietal Lobule.

### Section II: The comparison of different activation detection methods

#### 1. The results for activation detection with above-threshold event (Wang, et.al., 2019 ).

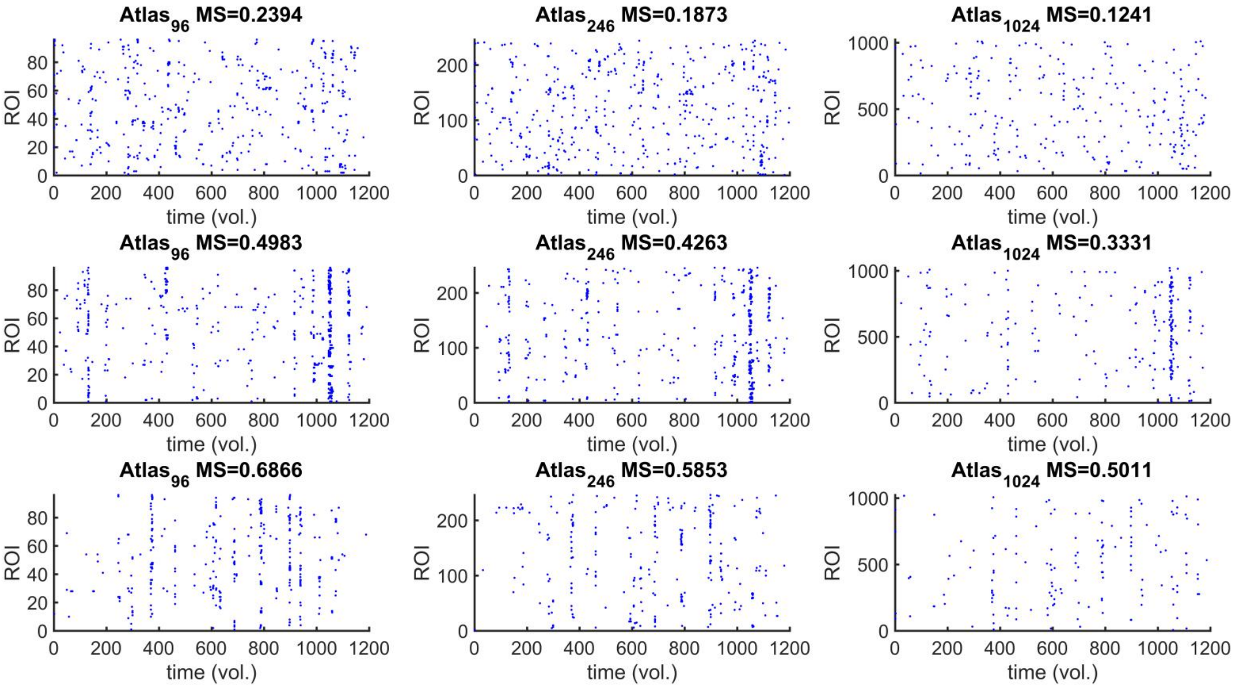


**Figure S15. Samples of above-threshold events for different level of MS obtained from atlas with different size.** The events were detected with threshold = 2.7 SD for $atlas_{96}$, 3 SD for $atlas_{246}$, and 3.6 SD for $atlas_{1024}$.

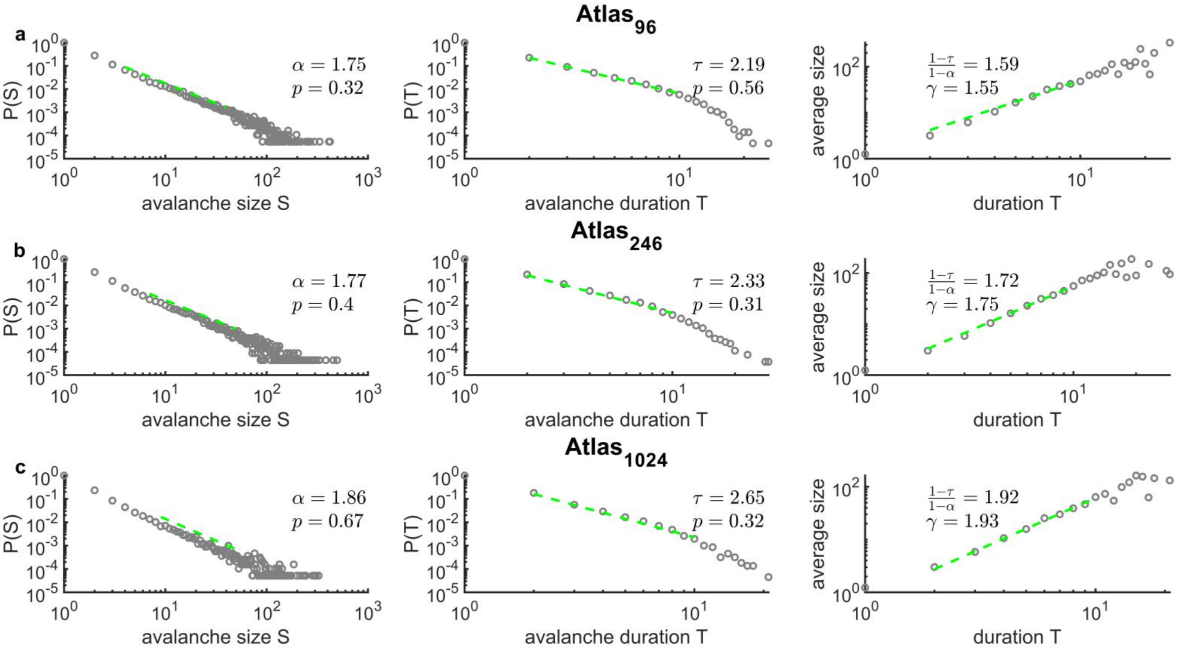


**Figure S16. Avalanche statistics obtained from group-level analysis using the above-threshold events.** The power-law distributions of avalanche sizes (left panel), avalanche durations (middle panel), and the scaling relation (right panel) obtained from $atlas_{96}$ (**a**), $atlas_{246}$ (**b**), and $atlas_{1024}$ (**c**). The events were detected with threshold = 2.7 SD for $atlas_{96}$, 3 SD for $atlas_{246}$, and 3.6 SD for $atlas_{1024}$. The values for power-law exponents of avalanche sizes $\alpha$ and avalanche durations $\tau$, as well as the Cluset's test results $p$ are indicated.

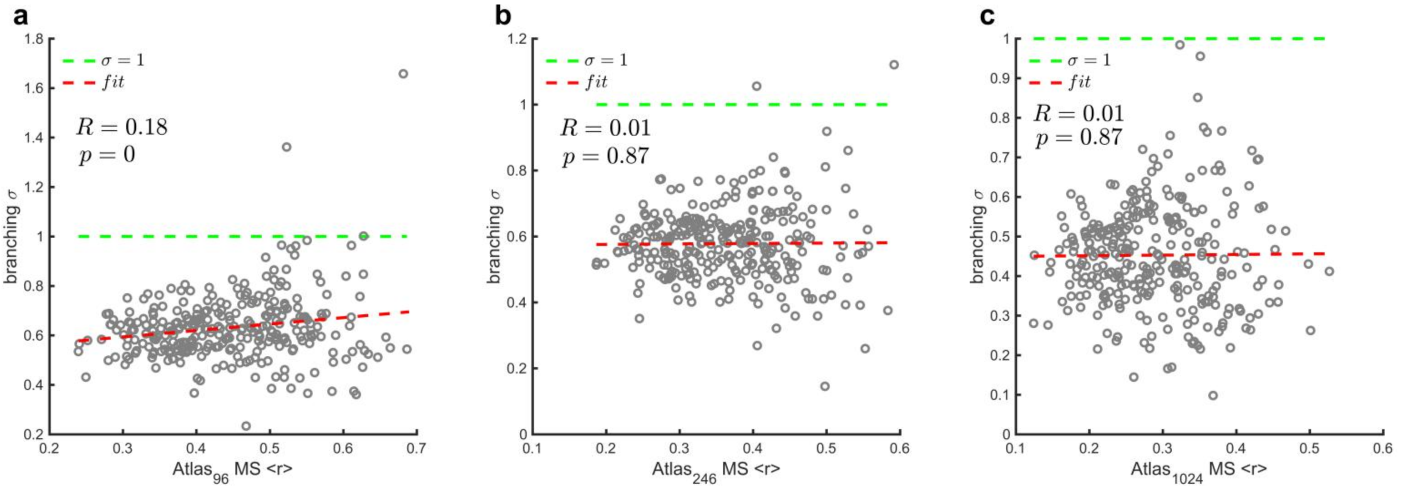


**Figure S17. The relationship between branching parameters and MS for above-threshold events.** The branching parameters $\sigma$ vs. MS $< r >$ for each subject obtained from atlas$_{96}$ (**a**), atlas$_{246}$ (**b**), and atlas$_{1024}$ (**c**). The green dashed lines indicate $\sigma = 1$. The Pearson correlation values $R$ and $p$ values are shown in the figure. The red dashed lines are the linear regression. The events were detected with threshold = 2.7 SD for atlas$_{96}$, and 3 SD for atlas$_{246}$, 3.6 SD for atlas$_{1024}$.

**2. The results for activation detection with threshold-crossing event (Bocaccio, et al., 2019).**

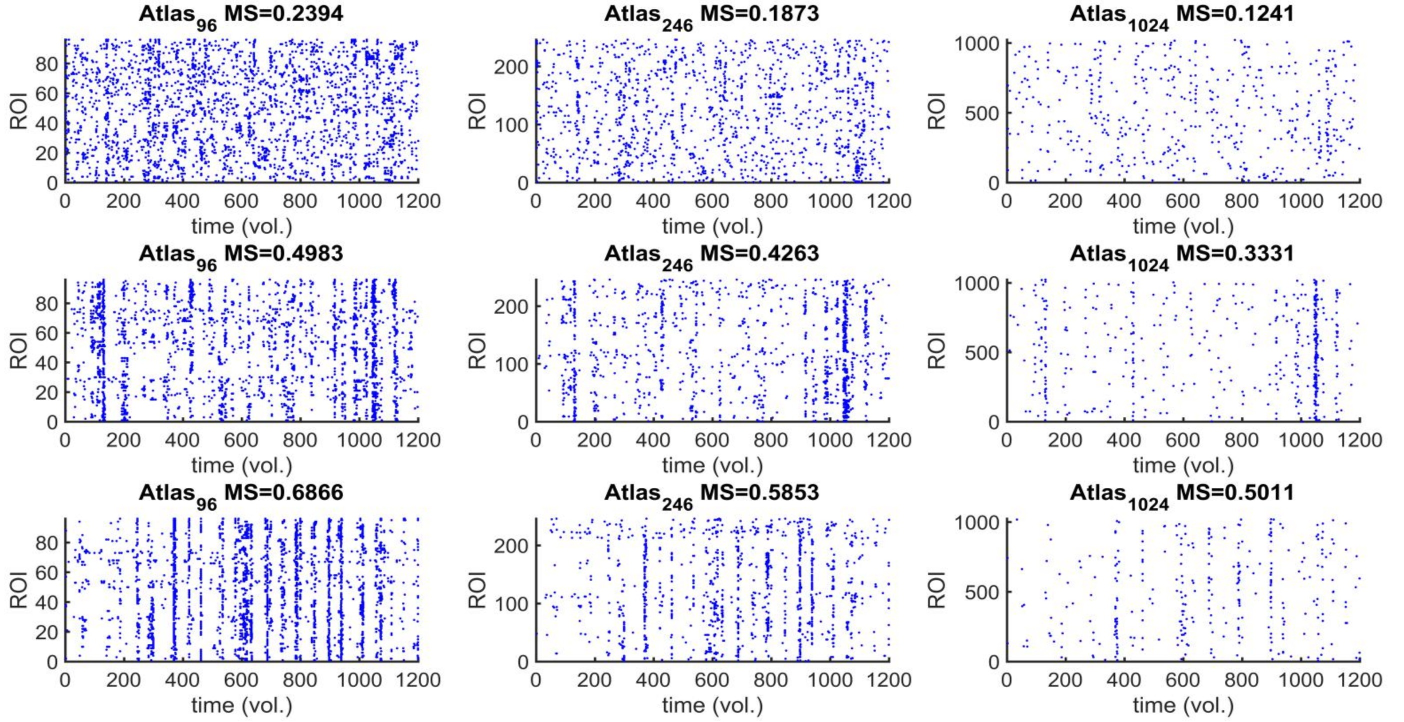


**Figure S18. Sample of threshold-crossing events for different levels of MS obtained with different atlas size.** The events were detected with threshold = 1.95 SD for atlas$_{96}$, 2.6 SD for atlas$_{246}$, and 3.4 SD for atlas$_{1024}$.

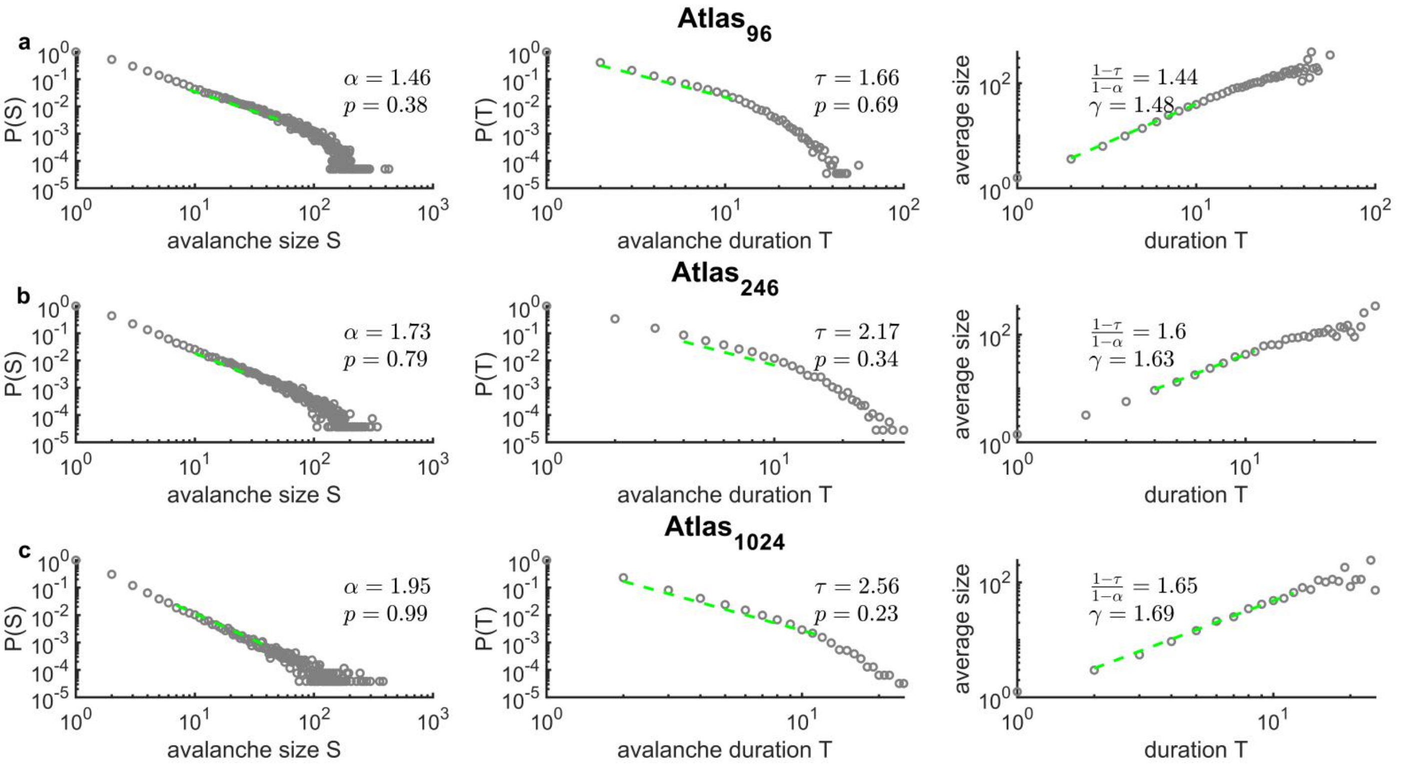


**Figure S19. Avalanche statistics obtained from group-level analysis using the threshold-crossing events.** The power-law distributions of avalanche sizes (left panel), avalanche durations (middle panel), and the scaling relation between avalanche sizes and durations (right panel) for $atlas_{96}$ **(a)**, $atlas_{246}$ **(b)**, and $atlas_{1024}$ **(c)**. The events were detected with threshold = 1.95 SD for $atlas_{96}$, 2.6 SD for $atlas_{246}$, and 3.4 SD for $atlas_{1024}$. The values for power-law exponents of avalanche sizes α and avalanche durations $\tau$, as well as the Cluset's test results $p$ are indicated.

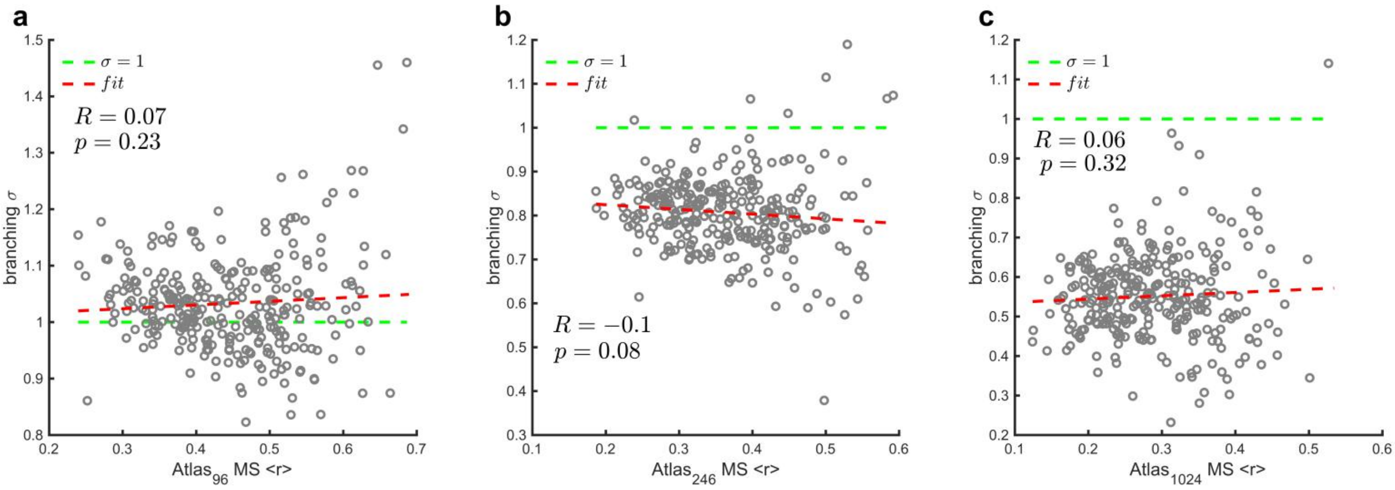


**Figure S20. The relationship between branching parameters and MS for threshold-crossing events.** The branching parameters $\sigma$ vs. MS $< r >$ for each subject for $atlas_{96}$ **(a)**, $atlas_{246}$ **(b)**, and $atlas_{1024}$ **(c)**. The green dashed lines indicate $\sigma = 1$. The Pearson correlation values $R$ and $p$ values are shown in the figure. The red dashed lines are the linear regression. The events were detected with threshold = 1.95 SD for $atlas_{96}$, 2.6 SD for $atlas_{246}$, and 3.4 SD for $atlas_{1024}$.
